# Bursting with potential: How sensorimotor beta bursts develop from infancy to adulthood

**DOI:** 10.1101/2023.05.09.539976

**Authors:** Holly Rayson, Maciej J Szul, Perla El-Khoueiry, Ranjan Debnath, Marine Gautier-Martins, Pier F Ferrari, Nathan Fox, James J Bonaiuto

## Abstract

Beta activity is thought to play a critical role in sensorimotor processes. However, little is known about how activity in this frequency band develops. Here, we investigated the developmental trajectory of sensorimotor beta activity from infancy to adulthood. We recorded electroencephalography (EEG) from adults, 12-month-olds, and 9-month-olds while they observed and executed grasping movements. We analysed ‘beta burst’ activity using a novel method that combines time-frequency decomposition and principal component analysis (PCA). We then examined the changes in burst rate and waveform motifs along the selected principal components. Our results reveal systematic changes in beta activity during action execution across development. We found a decrease in beta burst rate during movement execution in all age groups, with the greatest decrease observed in adults. Additionally, we identified four principal components that defined waveform motifs that systematically changed throughout the trial. We found that bursts with waveform shapes closer to the median waveform were not rate-modulated, whereas those with waveform shapes further from the median were differentially rate-modulated. Interestingly, the decrease in the rate of certain burst motifs occurred earlier during movement and was more lateralized in adults than in infants, suggesting that the rate modulation of specific types of beta bursts becomes increasingly refined with age.

## Introduction

Modulation of neural activity in different frequency bands is a classic signature of many sensory, motor, and cognitive processes. However, the generative mechanisms that drive this activity and the computational functions that band-specific neural activity subserves remain unclear. Developmental research provides a unique opportunity to elucidate these processes by allowing investigation into the unfolding relationship between frequency-specific neural activity and emerging abilities in various domains (Munakata et al., 2004). While most research with infants has focused on theta and alpha bands (Orekhova et al., 2006; Saby and Marshall, 2012; Schaworonkow and Voytek, 2021; Thorpe et al., 2016), very little is known about the development of neural activity in other frequency bands, especially the beta band (Cuevas et al., 2014; Perone and Gartstein, 2019), despite its crucial role in sensorimotor and cognitive control in adults (Järveläinen et al., 2004; Pfurtscheller and Neuper, 1997).

Traditionally, beta activity was thought to occur as a sustained oscillation with slow task-related amplitude modulations. In sensorimotor cortex, trial-averaged beta power decreases prior to movement and increases afterwards (Alayrangues et al., 2019; Cassim et al., 2001; Cheyne, 2013; Donner et al., 2009; Erbil and Ungan, 2007; Haegens et al., 2011; Keinrath et al., 2006; Kilavik et al., 2013; Kilner et al., 2003; Leocani and Comi, 2006; McFarland et al., 2000; Meirovitch et al., 2015; Miller et al., 2010; G. Pfurtscheller et al., 1996; Pfurtscheller and Lopes da Silva, 1999; Pogosyan et al., 2009; Salenius and Hari, 2003; Tan et al., 2016; Tzagarakis et al., 2010, 2015). This view assumes that single trial beta activity resembles trial-averaged activity, therefore, the pre-movement decrease and subsequent increase in beta power were named event-related desynchronisation (ERD) and synchronisation (Neuper et al., 2006; G Pfurtscheller et al., 1996; Pfurtscheller and Lopes da Silva, 1999). These terms imply that at rest, neuronal populations are synchronously activated at a frequency within the beta range, and that this synchrony is disrupted during motor preparation and movement, followed by a period of enhanced synchrony post-movement. However, it has now been consistently shown that at the single trial level, beta activity predominantly occurs as transient bursts rather than oscillations (Cagnan et al., 2019; Diesburg et al., 2021; Feingold et al., 2015; Haufler et al., 2022; Howe et al., 2011; Karvat et al., 2020; Kosciessa et al., 2020; Little et al., 2019; Lofredi et al., 2019; Sherman et al., 2016; Shin et al., 2017; Sporn et al., 2020; Torrecillos et al., 2018; Wessel, 2020), changes in burst probability closely track trial-averaged beta power (Feingold et al., 2015; Howe et al., 2011; Little et al., 2019; Rayson et al., 2022; Sherman et al., 2016), and the burst timing is highly predictive of behaviour (Diesburg et al., 2021; Echeverria-Altuna et al., 2021; Enz et al., 2021; Feingold et al., 2015; Haufler et al., 2022; Heideman et al., 2020; Howe et al., 2011; Karvat et al., 2020; Khawaldeh et al., 2020; Kosciessa et al., 2020; Law et al., 2022; Little et al., 2019; Sherman et al., 2016; Shin et al., 2017; Sporn et al., 2020; Torrecillos et al., 2018; Walsh et al., 2022; Wessel, 2020; West et al., 2022). This suggests that bursts are more informative about sensorimotor processing than beta power, and that averaging bandpass-filtered power over trials may obscure important features and functions of beta activity

Thus far, most investigations of beta bursts have treated them as homogeneous events, considering only the timing or rate of burst occurrence. However, recent research suggests that bursts can be highly diverse in terms of time-frequency-based features such as power, peak frequency, and frequency span (Duchet et al., 2021; Enz et al., 2021; Feingold et al., 2015; Szul et al., 2022; Zich et al., 2020), as well as in waveform shape in the temporal domain (Szul et al., 2022). Beta burst generation is proposed to involve temporally aligned synaptic inputs to deep and superficial layers of sensorimotor cortex (Bonaiuto et al., 2021; Sherman et al., 2016), and the timing, duration, and strength of these inputs can modulate their waveform shape and temporal characteristics (Szul et al., 2022). Differences in waveform shape, in turn, drive differences in time-frequency-based features (Jones, 2016; Szul et al., 2022). In adult sensorimotor cortex, beta bursts have a stereotyped mean waveform shape (Baker et al., 1997; Bonaiuto et al., 2021; Brady and Bardouille, 2022a; Cagnan et al., 2019; Cole et al., 2017; Karvat et al., 2020; Kosciessa et al., 2020; Sherman et al., 2016), but individual burst waveforms are highly diverse and occur within motifs which are differentially rate-modulated before, during, and after movements (Szul et al., 2022). It has been suggested that different types of beta bursts might therefore play different roles in sensorimotor processes (Szul et al., 2022), but it is not yet known what these processes are.

In terms of sensorimotor development, the first year of life is marked by rapid and significant changes in an infant’s perceptual and motor abilities (Adolph and Franchak, 2017; von Hofsten and Rosander, 2018), making this an ideal age range to probe the relationship between beta activity and movement. Very few EEG studies have looked at sensorimotor beta power in early development (Meyer et al., 2016, 2011; Niemarkt et al., 2011; Ogawa et al., 1984; Samson-Dollfus et al., 1983; van Elk et al., 2008), though studies with young children have demonstrated movement-related decreases in beta power (Bryant and Cuevas, 2019; Cheyne et al., 2014; Gaetz et al., 2010; Liao et al., 2015). Beta peak frequency increases with age in older children and adults (Johnson et al., 2019; Trevarrow et al., 2019), but infant studies that have looked at beta tend to use the canonical adult beta frequency band or one just above infant alpha. In one very recent study, we demonstrated that beta activity does indeed appear ‘bursty’ in 12-month-olds (Rayson et al., 2022), but to our knowledge no other study has looked at beta bursts in developmental populations. If different types of beta bursts are reflective of different sensorimotor processes, they may be differentially rate-modulated as these processes develop in infancy.

With the aim of using sensorimotor development to unveil the functional relevance of beta burst activity, here we analysed high-density electroencephalography (EEG) recordings obtained during execution and observation of a reaching-and-grasping action in 9- and 12-month-old infants, and an adult comparison group (Cannon et al., 2014; Yoo et al., 2016). We found that, like theta and alpha frequencies (Berchicci et al., 2011; Marshall et al., 2002; Orekhova et al., 2006), beta peak frequency increases over the first year of life and beyond. We demonstrate that, as in adults, beta activity in infants occurs as transient, rate-modulated bursts. Burst rate decreases around the onset of movement in adults, but not until the hand contacts the object in 9- and 12-month-olds. We use dynamic time warping to show that infant beta bursts have the same waveform shape as adult bursts, differing only in that they are stretched in time, thus explaining the difference between infants and adults in beta peak frequency. Principal component analysis of burst waveforms was then used to identify common waveform motifs between ages that are increasingly rate-modulated, earlier during the movement, and in an increasingly lateralized way, from infancy to adulthood. These results highlight the importance of considering the diverse features and potential functions of beta bursts, and suggest that developmental changes in burst rate and waveform motifs reflect the maturation of thalamic and cortical projections to sensorimotor cortex which shape the contralateral organisation of the motor system, and thereby support the emergence of internal models for motor preparation and planning.

## Materials and Methods

### Participants

Forty-four full-term 9-month-old infants (25 female, age range 8.6-9.93 months), 46 full-term 12-month-old infants (26 female, age range 11.2–12.93 months), and 23 adults (10 females, age range 18–22 years) were recruited for a study on the neural bases of action execution and observation. Sixteen 9-month-old infants were excluded due to unusable EEG data before preprocessing (*N* = 5), fussiness after applying the EEG electrode not (N = 6), and never grasping the toy during the experiment (*N* = 1). Thirteen 12-month-old infants were excluded due to unusable EEG data before preprocessing (*N* = 6), becoming distressed shortly after applying the EEG electrode net (*N* = 6), and recording failure (*N* = 1). One adult participant was excluded due to a data recording error. The final sample therefore included 28 9-month-old infants, 33 12-month-old infants, and 22 adults. All infants were typically developing with no known or suspected neurodevelopmental or medical diagnoses. Before an infant’s participation in the study, informed consent was obtained from the infant’s parents. All adults had normal or corrected to normal vision and did not have any neurological disorder. They provided informed consent before participating in the study. The experiment was approved by the University of Maryland Institutional Review Board.

### Procedure and task

Both infants and adult participants performed the same task. The infants were seated on their caregiver’s lap, whereas the adults were seated in a chair, approximately 40 cm away from a black puppet stage (99 cm × 61 cm × 84 cm), which was placed on a table. The area surrounding the stage was covered by black curtains to conceal the experimenter and any equipment from the participant’s view. A video camera was placed behind the presenter to record any significant behavioural events that occurred during the testing, and the caregivers of the infants were told to remain passive.

The task comprised two conditions: observation and execution. To start the observation trials, the curtain was raised, revealing a female presenter. The presenter made eye contact with the participant and then looked towards a toy that was positioned in the centre of the stage, but not within the participant’s reach. The presenter then picked up the toy using a hand-operated claw-like tool, brought it to themselves, and briefly shook it. The curtain was lowered to mark the end of this trial, which lasted approximately 4 seconds. For the execution trials, a toy was placed on the table, and while the presenter was hidden from the participant’s view, the table was pushed towards the participant within reach as the curtain was raised. Participants were given about 60 seconds to reach for the toy. The table was then pulled back and the curtain was lowered to mark the end of this trial. The order of the observation and execution trials was pseudo-randomized.

Ten different toys were utilised, with each toy being used in two observation and two execution trials. For adults, there was a maximum of 20 trials per condition per adult. On average, 9-month-old infants each completed 14.21 trials (*SD* = 4.82), 12-month-old infants completed 13.30 trials (*SD* = 4.82), and all adults completed 19.64 trials (*SD* = 1.71) per condition.

### Behavioural coding

Behavioural events were captured using video recorded at a resolution of 640 × 480 pixels and 30 Hz frame rate, and synchronised with the EEG recording. Two coders viewed the video offline and identified the frames when various events occurred (100 % overlap). For the execution condition, these events were the first touch of the toy with the hand, and the completion of the grasp. The same events were coded for the observation condition, but the first touch was the time when the presenter first touched the toy with the tool, and grasp completion was the completion of the grasp of the toy with the tool. For the execution condition, the hand or hands used to grasp the toy were also coded (in the observation condition the presenter always used their right hand). The coders achieved an inter-rater agreement of 84-94% within a three-frame window of approximately 100 ms (9m execution: first touch 93%, grasp complete 89%; 9m observation: first touch 92%, grasp complete 92%; 12m execution: first touch 93%, grasp complete 86%; 12m observation: first touch 91%, grasp complete 94%; adult execution: first touch 84%, grasp complete 94%; adult observation: first touch 93%, grasp complete 94%). These coded time points, as well as the start and end of the trials, were then used to segment the EEG data into epochs centred on these events.

### EEG recording and preprocessing

EEG was recorded using a 65-channel HydroCel Geodesic Sensor Net (Electrical Geodesics, Inc., Eugene, OR). The vertex (Cz) electrode was used as an online reference. EEG data were sampled at 500 Hz using EGI’s Net Station (v4.5.4) software. Impedances were kept below 100 kΩ. After recording, EEG data were exported to a MatLab compatible format using NetStation software for offline processing.

Both infant and adult datasets were preprocessed using Matlab R2018a with a custom version of the MADE pipeline (Debnath et al., 2020), which was modified to include artefact detection routines from the NEAR pipeline (Kumaravel et al., 2022). The data were high pass filtered at 1 Hz and low pass filtered at 100 Hz using EEGLAB v14.1.1 (Delorme and Makeig, 2004) FIR filters. Artefact-laden channels were identified and removed using the local outlier factor metric from the NEAR pipeline with an adaptive threshold starting at 2.5 for outlier detection (9m infants: 0–13 channels, *M* = 2, *SD* = 2.83; 12m infants: 0–8 channels, *M* = 1.58, *SD* = 1.85; adults: 0–5 channels, *M* = 1.18, *SD* = 1.59). Artefact subspace reconstruction (ASR) from the NEAR pipeline was then used to identify and correct non-stereotyped artefacts, using a cut-off parameter, k = 13. Stereotyped artefacts were then detected and removed using the independent component analysis (ICA) -based techniques from the MADE pipeline. ICA was performed on an identical copy of the dataset, which was first segmented into 1 s epochs. Noisy epochs in the copied dataset were removed using a voltage threshold of ± 1000 μV. After ICA decomposition, independent components (ICs) were transferred from the copied dataset to the original dataset, which was used from then on. Artifactual ICs were removed from the original dataset using the EEGLAB Adjusted-ADJUST plugin (Leach et al., 2020; Mognon et al., 2011; 9m infants: 3–50 components, *M* = 18.14, *SD* = 11.67; 12m infants: 6-52 components, *M* = 18.48, *SD* = 10.22; adults: 6–34 components, *M* = 16.77, *SD* = 6.98). The data were then divided into 3.6 second long epochs, centred on four events: the start of the trial, the moment the toy was first touched, the completion of the grasp, and the end of the trial. A voltage threshold of + /− 150 µV was used to detect artefacts in each channel during each epoch. For each epoch, if more than 10% of the channels contained artefacts, the epoch was removed (9m infants: 0-30 epochs, *M* = 5.68, *SD* = 7.95; 12m infants: 0-50 epochs, *M* = 5.94, *SD* = 11.79; adults: no epochs removed), otherwise the artefacted channels were removed and interpolated for that epoch. After artefact rejection, any remaining missing channels were interpolated, and the data were average re-referenced. Finally, line noise (60 Hz) was removed using an iterative version of the Zapline algorithm (de Cheveigné, 2020) implemented in the MEEGKit package (https://nbara.github.io/python-meegkit/), using a window size of 10 Hz for polynomial fitting and 2.5 Hz for noise peak removal and interpolation. For the execution condition, epochs during trials in which no grasp was made, the grasp used two hands, the toy was touched twice before grasping, or the grasp took longer than 1.6 seconds to complete were rejected. Following preprocessing, for each epoch, all participants with at least 5 trials were included in the following analyses (Table S1). All source code for preprocessing is available at https://github.com/danclab/dev_beta_claw/tree/main/preprocessing.

### Kinematics analysis

Before excluding trials in which two hands were used to grasp the toy, the type of grasp (unimanual or bimanual) was analysed for the execution condition using a generalised linear mixed model with a binomial distribution and logit link function with age as a fixed effect and subject-specific intercepts as random effects. The hand used (left or right) for unimanual movements in the execution was then analysed in the same way. These were not analysed for the observation condition because the adult actor always performed the action using their right hand. The time from the start of the trial until the first touch of the toy (reach duration), from the first touch until grasp completion (grasp duration), and from grasp completion until the end of the trial (manipulation duration), were analysed separately for the execution (including only unimanual trials) and observation conditions using linear mixed models including age as a fixed effect, and subject-specific intercepts as random effects. All analyses of movement kinematics were conducted using R (v3.6.1; R Core Team, 2022) and lme4 (v1.1.29; Bates et al., 2014). Fixed effects were assessed using type II Wald *Χ*^2^ tests (car v3.1.0; Fox et al., 2019). Pairwise Tukey-corrected follow-up tests were run using estimated marginal means from the emmeans package (v1.7.3; Lenth et al., 2020).

### Burst analyses

Power spectral densities (PSDs) were computed from 0.1 to 100 Hz with the MNE-Python toolbox (Gramfort et al., 2014) using Welch’s method (Welch, 1967) with a window size of 1 s, 50% overlap, and 10 times oversampling, resulting in a frequency resolution of 0.1 Hz. For each participant (infants and adults), this was applied to all data (i.e. from both epochs of the observation and execution conditions). For each channel, the power spectral density was parameterized using specparam (Donoghue et al., 2020) to obtain estimates of the aperiodic and periodic spectral components. The periodic spectra were then averaged over all electrodes in the C3 and C4 clusters (E16, E20, E21, E22, E41, E49, E50, E51), and then across participants within each age group. Group-level spectral peaks in the periodic component were identified using an iterative procedure in which a Gaussian function was fitted to the global maximum and subtracted from the periodic spectral density. The procedure was then repeated using the result of the subtraction, continuing until there are no more peaks above the noise floor (1 standard deviation across all frequencies). For each peak, the frequency band limits were determined by computing the full width half maximum (FWHM) of the peak power, and only bands below 10 Hz with a FWHM of at least 1 Hz or above 10 Hz with a FWHM of at least 3 Hz were retained.

Lagged coherence was computed for all data from each participant (e.g. from both the observation and execution conditions) from 5 to 100 Hz in 1 Hz increments and 2–4.5 cycles in increments of 0.1 cycles (Fransen et al., 2015) using FieldTrip v20190329 (Oostenveld et al., 2011). We used overlapping epochs with lag- and frequency-dependent widths. Fourier coefficients were obtained for each epoch using a Hann-windowed Fourier transform. Because lagged coherence depends on data SNR (Fransen et al., 2015), we normalised lagged coherence values for each participant by the maximum lagged coherence over all channels, frequencies, and lags for that participant.

We used the superlet transform (Moca et al., 2021) to compute single trial TF decompositions for each electrode with optimally balanced time and frequency resolution. We used an adaptive superlet transform based on Morlet wavelets with varying central frequency (1 - 100 Hz) and number of cycles (4 cycles) under a Gaussian envelope. The order was linearly varied from 1 to 40 over the frequency range.

For each channel of the C3 and C4 clusters, within the beta frequency band identified, we used an adaptive burst detection algorithm to detect all potentially relevant burst events across a wide range of beta amplitudes during the execution condition (Szul et al., 2022). Similar to the method used for peak detection with the PSDs, the algorithm first subtracts the estimated aperiodic spectrum from each single trial TF decomposition and on each iteration, detects the global maximum amplitude in TF space and fits a two-dimensional Gaussian to this peak by computing the symmetric full width at half maximum (FWHM) in the time and frequency dimensions. The Gaussian is then subtracted from the TF decomposition, and the next iteration operates on the resulting residual TF matrix. This process continues until there are no global maxima above the noise floor remaining (2 standard deviations above the mean amplitude over all time and frequency bins, recomputed on each iteration). We applied the algorithm to TF data within a window 5 Hz wider than the identified beta band on either side, but only bursts with a peak frequency within the band were retained for further analysis.

The waveform for each detected burst was extracted from the "raw" time series (only 1 Hz high pass and 100 Hz low pass filtered during preprocessing), based on their peak time. To eliminate the influence of slower event-related potential (ERP) dynamics on burst waveforms, the epochs were first averaged in the temporal domain to calculate the ERP, and this was then removed from the signal for each trial. The width of the time window for waveform extraction was determined using the FWHM of lagged coherence averaged within the beta band. The time series within this window, centred on the peak time, was then extracted from the trial time series. To find the signal deflection corresponding to the peak in TF amplitude, the burst waveforms were aligned by band pass filtering them within their detected frequency span (using a zero-phase FIR filter with a Hamming window), calculating their instantaneous phase using the Hilbert transform, and re-centering the "raw" waveform (before band pass filtering) around the phase minimum closest to the peak time detected in TF space (Boto et al., 2022; Szul et al., 2022). If this time point was more than 30 ms away from the TF-detected peak time, the burst was discarded. The DC offset was then subtracted from the resulting waveform. Finally, to account for uncertainty in the orientation and source location of the dipoles that generated the signals measured by the EEG electrodes, the sign of burst waveforms in which the central deflection was positive was reversed (Jones et al., 2009; Szul et al., 2022). Open-source code for the burst detection algorithm can be found at https://github.com/danclab/burst_detection.

Because infants tended to use either hand to grasp the toy, the time courses of burst rate and beta power in the execution condition were analysed according to which electrode cluster (C3 or C4) was contralateral or ipsilateral to the hand used. In the observation condition, bursts were simply categorised according to the electrode cluster they were identified in (C3 or C4). The burst rate was computed by binning bursts in 25 ms time bins, and then smoothing the resulting histogram with a Gaussian kernel (width = 3 time points). Both burst rate and mean beta amplitude (averaged within the identified beta band) were baseline-corrected using the mean rate or amplitude during the 1.5 seconds before the start of the trial, and expressed as a percentage change from baseline. To identify significant deviations from the baseline, we used a one-sample cluster permutation test. The family-wise error rate (FWER) was controlled using a non-parametric resampling test with a maximum statistic (taken across all data points). A t-test with relative variance regularisation (“hat” adjustment; Ridgway et al., 2012; sigma = 0.001) was used as the statistic to minimise the impact of low variance data points and prevent spurious results. Threshold-free cluster enhancement (TFCE) was used to enhance the statistical power of cluster detection by utilising an adaptive threshold on the level of a single data point (starting threshold = 0, step = 0.01; Smith and Nichols, 2009).

Burst peak amplitude, peak frequency, frequency span, and duration were analysed using R (v3.6.1; R Core Team, 2022) with linear mixed models including age, or age, cluster, and their interactions as fixed effects, and subject-specific intercepts as random effects (lme4 v1.1.29; Bates et al., 2014). Because the peak frequency was necessarily different across ages due to the beta band identification procedure, the analysis of the effect of cluster on peak frequency included subject nested within age as random effects. Fixed effects were assessed using type III Wald *Χ^2^* tests (car v3.1.0; Fox et al., 2019). Pairwise Tukey-corrected follow-up tests were run using estimated marginal means from the emmeans package (v1.7.3; Lenth et al., 2020).

Burst waveforms from 9-month and 12-month-old infants were separately warped to adult burst waveforms using dynamic time warping (Giorgino, 2009). To account for differences in burst amplitude, we first normalised the median burst waveform for each group, and performed dynamic time warping on the normalised median infant burst waveforms using Rabiner-Juang step patterns (type 5c; Rabiner and Juang, 1993), with the normalised median adult burst waveform as the reference. The resulting alignments were then used to warp all infant bust waveforms.

To classify burst waveform shapes, principal component analysis (PCA, 20 components, implemented in the scikit-learn library; Pedregosa et al., 2011) was applied to the warped burst waveforms from all age groups (Szul et al., 2022). All detected bursts were then projected onto each principal component (PC), with each burst thus having a score for each component representing the shape of its waveform along that dimension. To determine which components were not simply caused by noisy signal fluctuations, we used a permutation approach (Vieira, 2012) to remove correlations between features (waveform time points). The matrix containing all burst waveforms was shuffled within each time point (column) independently, and PCA was applied to the shuffled matrix. The *p* value for each PC was then given by the probability of the proportion of variance explained being lower after shuffling than that for the unshuffled data. One hundred permutations were run for each component, with an alpha threshold of *p* = 0.0035, using Bonferroni correction for multiple comparisons. To evaluate consistency in PCA results across age groups, we ran a separate PCA for each age group, using only bursts detected from participants in that group. For each group PCA, we examined correlations between its eigenvectors and those of the global PCA by constructing a correlation matrix comparing each component from one PCA to each component of the other. Because the direction of each dimension identified by PCA is arbitrary, we took the absolute value of the correlation coefficient. To account for different potential ordering of components, we looked at the maximum correlation in each row of the matrix (i.e. for each component of the global PCA, the component of the group PCA most correlated with it). This same procedure was used to examine correlations between burst scores after projecting their waveforms onto each dimension of the global and group PCAs.

We proceeded to select principal components (PCs) that define dimensions along which the mean burst waveform shape varied during either the action observation or execution conditions. For each PC, the mean burst waveform score was computed for each epoch in 50 ms bins over both electrode clusters. Subsequently, the scores were smoothed using a Gaussian kernel with a width of 2 time points and baseline-corrected using the 1.5 seconds preceding the beginning of the trial. To ascertain when the mean score deviated from the baseline for each PC, a one-sample cluster permutation test was employed over all subjects from all age groups. To control the family-wise error rate (FWER), a non-parametric resampling test with a maximum statistic (taken across all data points) was used. A t-test with a relative variance regularisation (i.e., "hat" adjustment) was employed, and the threshold for significance was set at p = 0.000025, Bonferroni adjusted for multiple comparisons. TFCE was used, starting with a threshold of 0 and a step size of 0.2. Components with mean scores that significantly deviated from baseline during the toy touch, grasp completion, and trial end epochs in either the action observation or execution conditions were selected for further analysis.

To analyse the burst rate according to waveform shape, we binned bursts according to their component score, indicating their waveform shape (four quartiles), and the time during the trial in which they occurred (50 ms bins). For each component score quartile, we then baseline-corrected the burst rate as described above, and smoothed it using a Gaussian kernel (width = 2 time points). We used one-sample cluster permutation tests to determine when burst rates in each component quartile significantly deviated from the baseline. The family-wise error rate (FWER) was controlled using a non-parametric resampling test with a maximum statistic (taken across all data points). A t-test with a relative variance regularisation (“hat” adjustment; threshold p = 0.0000156, Bonferroni adjusted for multiple comparisons) and TFCE (starting threshold = 0, step = 0.2) was used.

All source code for these analyses is available at https://github.com/danclab/dev_beta_claw.

## Results

### Behavioural kinematics

Overall, the kinematics analysis revealed several significant differences in the behaviour of participants of different age groups. The great majority of movements in the execution condition were unimanual, and there was no difference in the amount of bimanual movements between age groups (*Χ*^2^(2) = 5.57, *p* = 0.062; 9m *M* = 0, *SD* = 0 % of trials; 12m *M* = 7.53, *SD* = 26.42 % of trials; adult *M* = 1.40, *SD* = 11.74 % of trials). There was a difference in the hand used for unimanual movements (*Χ*^2^(2) = 21.06, *p* < 0.001), with adults using the right hand (*M* = 84.62, *SD* = 36.12 % of trials) more than 9-month (*Z* = 4.00, *p* < 0.001; 9m *M* = 60.33, *SD* = 48.99 % of trials) and 12-month-olds (*Z* = 4.37, *p* < 0.001; 12m *M* = 60.15, *SD* = 49.02 % of trials), but there was no difference between 9- and 12-month-olds (*Z* = -0.35, *p* = 0.935).

In the execution condition, the time until the first touch of the toy was significantly different between the groups (*Χ*^2^(2) = 25.15, *p* < 0.001), with adults reaching the toy faster (*M* = 1.71, *SD* = 0.84 s) than both 9-month-olds (*t*(72.6) = -3.55, *p* = 0.002; 9m *M* = 8.00, *SD* = 13.97 s) and 12-month-olds (*t*(74.1) = -4.90, *p* < 0.001; 12m *M* = 9.63, *SD* = 16.81 s). There was no significant difference in the time to reach between 9- and 12-month-olds (*t*(84.8) = 1.30, *p* = 0.401). The duration of the grasping (*Χ*^2^(2) = 232.16, *p* < 0.001) and manipulation movements (*Χ*^2^(2) = 30.65, *p* < 0.001) were also significantly different between the groups, with adults performing faster grasping (*M* = 0.25, *SD* = 0.16 s) and manipulation movements (*M* = 1.41, *SD* = 0.40 s) faster than both 9-month-olds (grasping: *t*(69.9) = -13.74, *p* < 0.001; 9m *M* = 0.73, *SD* = 0.33 s; manipulation: *t*(67.6) = -5.19, *p* < 0.001; 9m *M* = 2.49, *SD* = 3.28 s) and 12-month-olds (grasping: *t*(71.7) = -12.81, *p* < 0.001; 12m *M* = 0.68, *SD* = 0.35 s; manipulation: *t*(69.5) = -4.29, *p* < 0.001; 12m *M* = 2.33, *SD* = 1.96 s). However, there was no significant difference in grasp duration (*t*(86.7) = -1.29, *p* = 0.403) or manipulation time (*t*(88.7) = -1.01, *p* = 0.574) between 9- and 12-month-olds. In summary, there were no differences in the kinematics of performed actions between 9 and 12 months, but adults performed faster reach, grasp, and manipulations movements than both infant age groups.

During the observation task, there was no significant difference between age groups in the duration of the observed reach (*Χ*^2^(2) = 0.49, *p* = 0.782; 9m *M* = 1.89, *SD* = 2.34 s; 12m *M* = 1.87, *SD* = 0.75 s; adult *M* = 1.81, *SD* = 0.92 s). However, the durations of the observed grasp (*Χ*^2^(2) = 11.83, *p* = 0.003) and manipulation movements (*Χ*^2^(2) = 7.45, *p* = 0.024) were significantly different, with 12-month-olds observing longer grasps (*t*(77.4) = -3.33, *p* = 0.004; 12m *M* = 0.68, *SD* = 0.31 s; adult *M* = 0.53, *SD* = 0.28 s) and manipulation movements (*t*(79.5) = 2.57, *p* = 0.032; 12m *M* = 2.76, *SD* = 0.38 s; adult *M* = 2.55, *SD* = 0.33 s). However, there were no significant differences in grasp or manipulation duration during observation between 9-month-olds and either 12-month-olds (grasp: *t*(82.0) = 2.23, *p* = 0.071; 9m *M* = 0.58, *SD* = 0.29 s; manipulation: *t*(80.4) = 0.28, *p* = 0.958; 9m *M* = 2.77, *SD* = 0.35 s) or adults (grasp: *t*(76.6) = -1.18, *p* = 0.472; manipulation: *t*(79.3) = -2.23, *p* = 0.072). There were therefore no consistent differences in the observed action kinematics with age.

### Sensorimotor beta occurs as transient bursts in infancy and adulthood

We first sought to confirm whether sensorimotor beta activity is ‘bursty’ rather than oscillatory in infants and adults. With this aim, we analysed EEG data recorded from clusters of electrodes centred around C3 and C4, while 9-month-old, 12-month-old, and adult participants reached for, grasped, and shook a toy, or observed an adult performing the same series of actions. After decomposing the power spectrum within these clusters into aperiodic and periodic components (Donoghue et al., 2020; Ostlund et al., 2022), we identified group-level periodic peaks within the canonical beta frequency range (13-30 Hz) in each age group from the periodic spectra averaged over all electrodes, conditions, and epochs. The identified beta frequency bands increased in peak frequency and range from 9 months to adulthood (9 months: 12.75 - 16.25 Hz, 12 months: 13.5 - 17.0 Hz, adults: 18.25 - 24.75 Hz, Figure 1a,d,g; Meyer et al., 2016; Rayson et al., 2022). PSDs and decomposed aperiodic and periodic spectra for each participant are shown in Figure S1.

**Figure 1.**
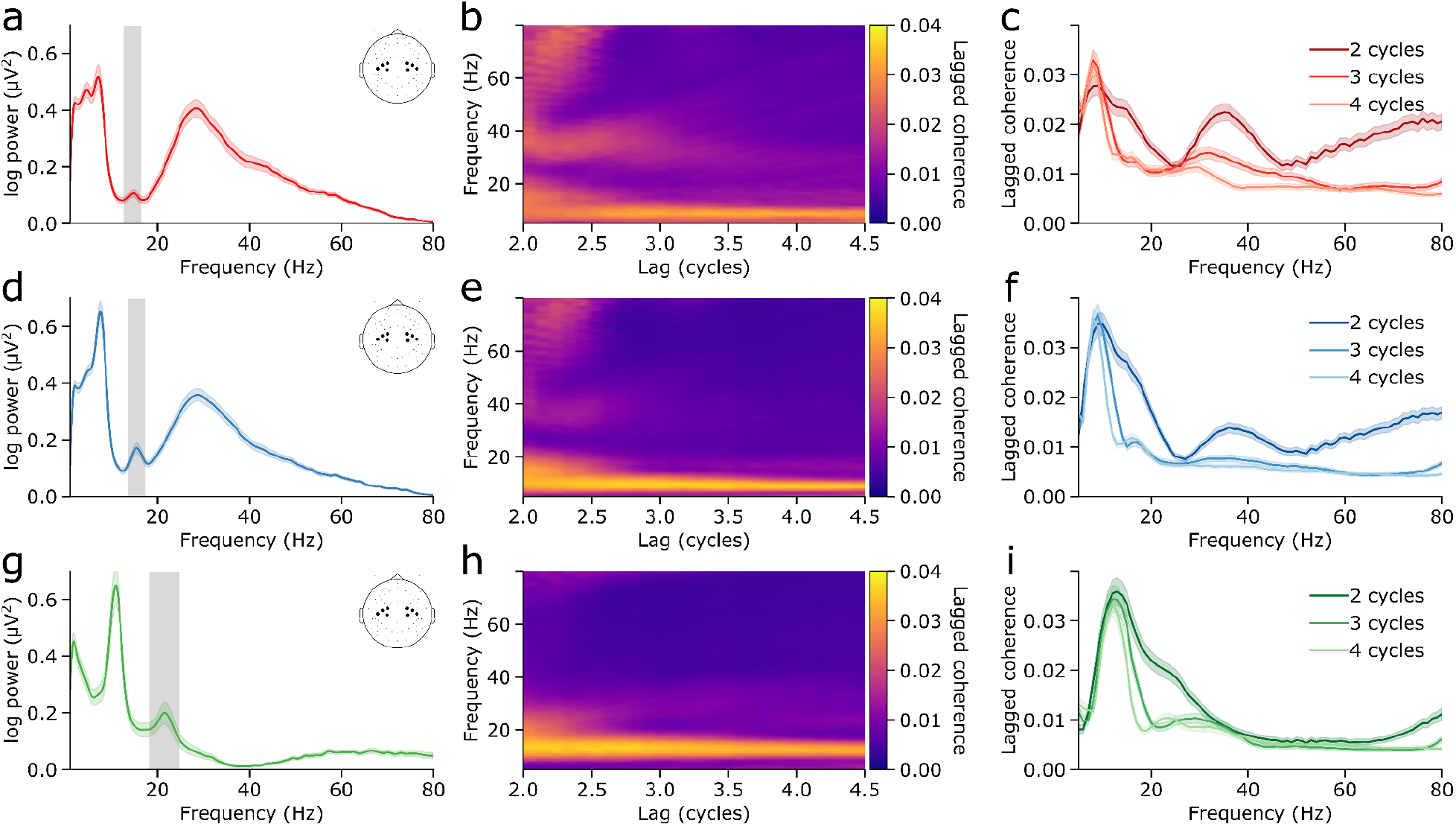
Beta peak frequency increases with age and beta activity consists of transient burst events in infancy and adulthood. a) Periodic power spectral density in the combined C3 and C4 cluster of the 9-month-old participants. The mean over participants is represented by the dark lines, and the standard error is represented by the shaded areas. The grey shaded region indicates the limits of the identified beta band, and the inset shows the electrodes included in the analysis. b) Mean lagged coherence in the combined C3 and C4 cluster across all 9-month-old participants for a range of frequencies and lags. There is high lagged coherence in the alpha band over a wide range of lags, but beta lagged coherence rapidly decreases after 2 cycles. C) Mean lagged coherence in the combined C3 and C4 cluster across all 9-month-old participants for lags of 2, 3, and 4 cycles. The mean is represented by the solid lines, and the standard error is represented by the shaded areas. A prominent peak appears in the alpha range at 2-4 cycles, while a beta peak is visible only at 2 cycles. d-f) As in a-c, for 12-month-old participants. g-i) As in a-c, for adult participants.

We evaluated the rhythmicity of activity across the entire frequency spectrum using lagged coherence from 2 to 4.5 cycles in increments of 0.1 cycles. Activity in the alpha/mu frequency range had a high lagged coherence value that was sustained over at least 4.5 cycles, but lagged coherence in the beta frequency bands fell rapidly after 2 cycles (Figure 1b,e,h), meaning that its phase was unpredictable after 2 cycles in the future. This indicates that sensorimotor alpha/mu activity occurred as a rhythmic oscillation sustained over several cycles, whereas beta activity occurred as transient bursts (Figure 1c,f,i).

Beta power and lagged coherence had different spatial topographies across age groups. Beta power localised to peripheral locations in 9-month and 12-month-old infants (Figure 2a,c), but to the C3 and C4 clusters and central frontal electrodes in adults (Figure 2e). However, in all age groups, lagged coherence in the beta band localised to the C3 and C4 electrodes at 2 lag cycles, and rapidly decreased with increasing cycles (Figure 2b,d,f). Mu/alpha and gamma activity had very different patterns of power and lagged coherence topographies (Figure S2). The beta band therefore has distinct spectral and spatial signatures in infancy and adulthood, and the spatial topography of beta power and lagged coherence confirmed our choice of electrode clusters.

**Figure 2.**
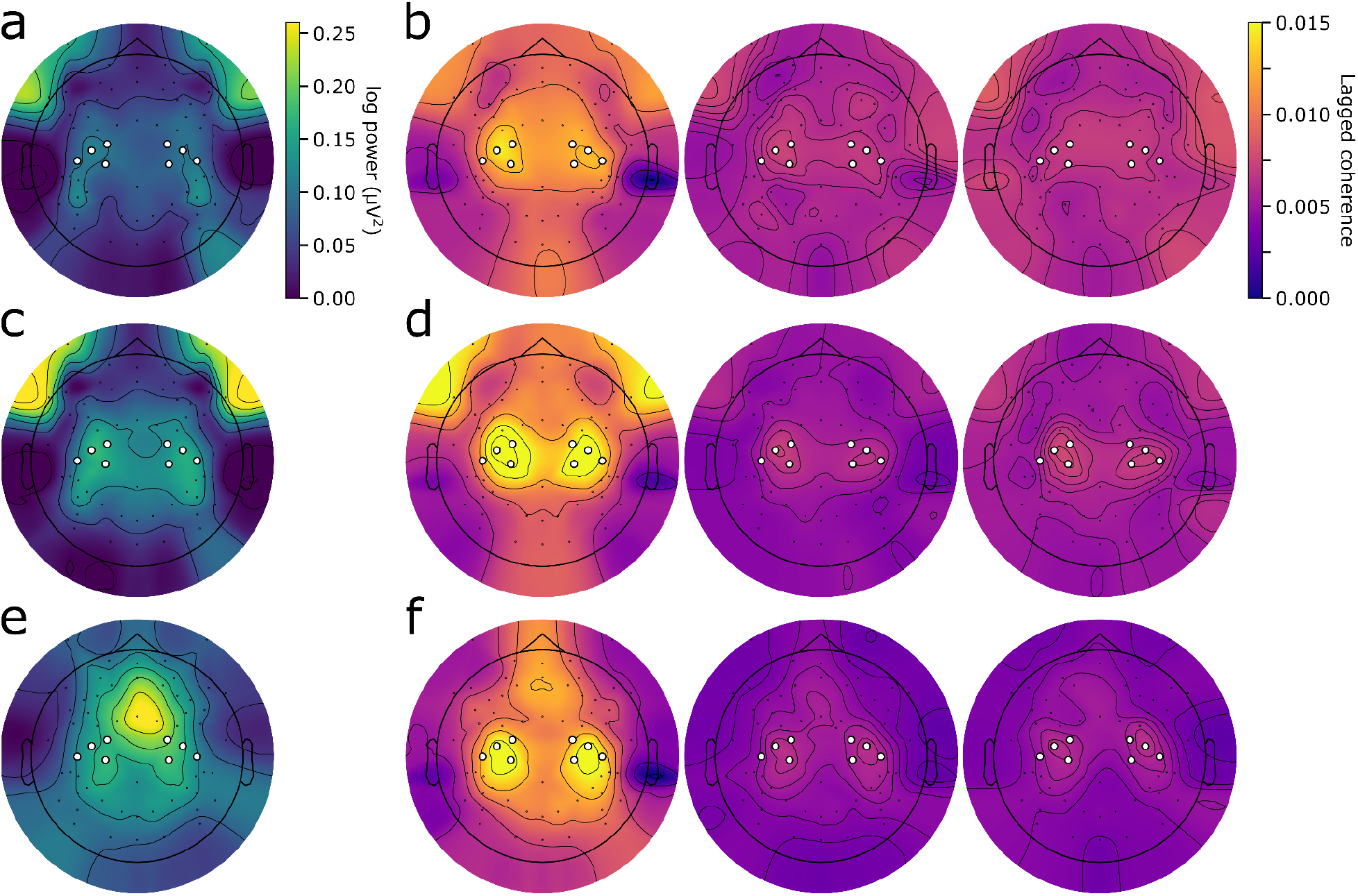
Power and lagged coherence localise beta to the C3 and C4 clusters. a) Topography of beta band periodic power (after subtraction of the aperiodic spectral density), averaged over 9-month-old participants. The white circles indicate electrodes included in the C3 and C4 clusters. Power in the beta band is most prominent in peripheral electrodes. b) Beta lagged coherence topographies at (from left to right) 2, 3, and 4 cycles averaged over 9-month-old participants. Lagged coherence in the beta band localises to the C3 and C4 electrodes and decreases rapidly after two cycles. c-d) As in a-b, for the 12-month-old participants. e-f) As in a-b, for the adult participants.

### Beta bursts have diverse time-frequency -based features

To detect beta bursts, we employed an adaptive, trial-based method to identify all potential burst events within each beta frequency range for each age group (Szul et al., 2022). We then compared time-frequency -based features of these bursts between age groups. Bursts across all age groups exhibited a wide range of peak amplitudes (Figure 3a), frequencies (Figure 3b), frequency spans (Figure 3c), and durations (Figure 3d). There was a main effect of age for burst peak amplitude (*Χ*^2^(2) = 118.75, *p* < 0.001), frequency span (*Χ*^2^(2) = 31.19, *p* < 0.001), and duration (*Χ*^2^(2) = 25.41, *p* < 0.001). There was no difference in peak amplitude between 9- and 12- month-olds (*Z* = -0.21, *p* = 0.975; 9m *M* = 0.97, *SD* = 0.63 μV; 12m *M* = 0.99, *SD* = 0.66 μV), but adult bursts had lower peak amplitudes than those detected in 9-month-old (*Z* = -9.61, *p* < 0.001; adult *M* = 0.42, *SD* = 0.30 μV) and 12-month-old infants (*Z* = -9.75, *p* < 0.001). Bursts detected in 12-month-olds had a slightly narrower frequency span as those detected in 9-month-olds (*Z* = -2.35, *p* = 0.049; 9m: *M* = 2.12, *SD* = 0.82 Hz; 12m: *M* = 2.08, *SD* = 0.82 Hz), and adult bursts were narrower than those of 9-month (*Z* = -5.57, *p* < 0.001; adult: *M* = 2.00, *SD* = 0.82 Hz) and 12-month-olds (*Z* = -3.56, *p* = 0.001). Finally, beta bursts were similar in duration at 12 months compared to at 9 months (*Z* = 0.13, *p* = 0.990; 9m: *M* = 4.42, *SD* = 2.03 cycles; 12m: *M* = 4.45, *SD* = 2.19 cycles), but lasted more cycles in adults compared to 9- (*Z* = 4.46, *p* < 0.001; adult: *M* = 4.99, *SD* = 2.80 cycles) and 12-month-olds (*Z* = 4.49, *p* < 0.001). In summary, while bursts did not differ in peak amplitude or duration between 9- and 12-month-olds, they decreased in amplitude, decreased in frequency span, and increased in duration from infancy to adulthood.

**Figure 3.**
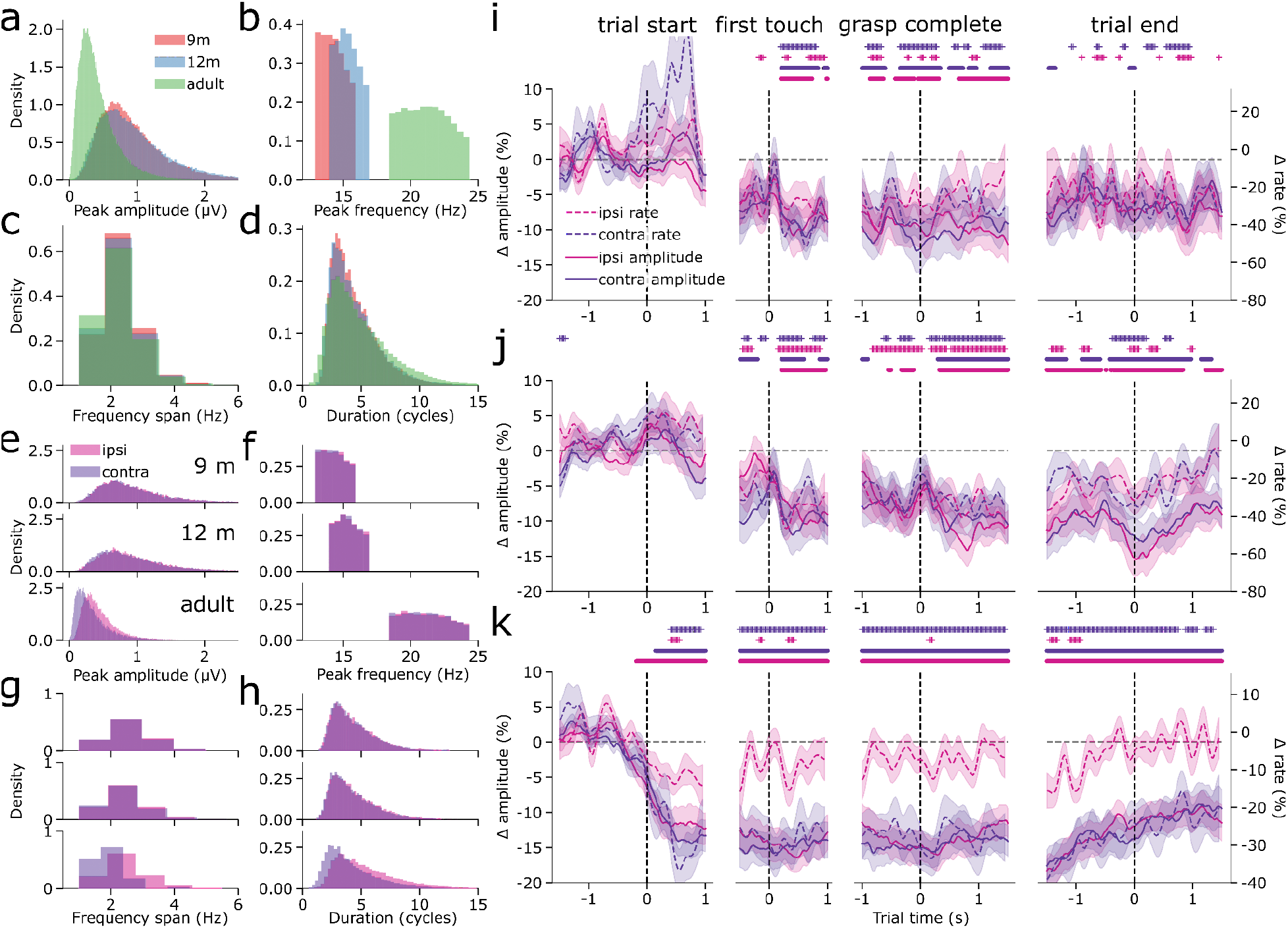
Beta bursts are diverse in infancy and adulthood. a-d) The distributions of beta burst peak amplitude (a), peak frequency (b), frequency span (c), and duration (d) for the 9-month-old, 12-month-old, and adult participants over conditions, epochs, and clusters. e-h) The distributions of burst peak amplitude (e), peak frequency (f), frequency span (g), and duration (h) for the (rows, from top to bottom), 9-month-old, 12-month-old, and adult participants for the ipsilateral and contralateral electrode clusters. i) The mean burst rate (dashed lines; shaded area shows the standard error) and mean beta amplitude (solid lines; shaded area shows the standard error) in the ipsilateral and contralateral clusters for the 9-month-old participants in the execution condition. The coloured dots and asterisks indicate times in which beta amplitude or burst rate significantly deviated from baseline. j) As in i, for the 12-month-old participants. k) As in i, for the adult participants.

We then sought to determine if there was any movement-related lateralisation in burst TF-based features and how this might change with age (Figure 3e-h). We categorised bursts in the execution condition based on whether the electrode cluster they were detected from was contra- or ipsilateral to the performed movement. Our results revealed an age-cluster interaction for burst peak amplitude (*Χ*^2^(2) = 379.91, *p* < 0.001), peak frequency (after accounting for age-specific frequency ranges; *Χ*^2^(2) = 12.54, *p* = 0.002), frequency span (*Χ*^2^(2) = 1,203.56, *p* < 0.001), and duration (*Χ*^2^(2) = 2184.24, *p* < 0.001). At 9 months, there was no difference in burst peak amplitude between hemispheres (*Z* = -0.09, *p* = 0.927; contralateral: *M* = 0.98, *SD* = 0.67 μV; ipsilateral: *M* = 0.99, *SD* = 0.63 μV). However, at 12 months and in adults, contralateral bursts had lower amplitudes than ipsilateral bursts (12 months: *Z* = -4.92, *p* < 0.001; contralateral: *M* = 0.97, *SD* = 0.72 μV; ipsilateral: *M* = 1.00, *SD* = 0.66 μV; adults: *Z* = -35.31, *p* < 0.001; contralateral: *M* = 0.35, *SD* = 0.29 μV; ipsilateral: *M* = 0.44, *SD* = 0.27 μV). Contralateral and ipsilateral bursts had the same peak frequency at 9 months (*Z* = -1.23, *p* = 0.217; contralateral: *M* = 14.31, *SD* = 0.97 Hz; ipsilateral: *M* = 14.33, *SD* = 0.97 Hz) and 12 months (*Z* = -0.50, *p* = 0.616; contralateral: *M* = 15.32, *SD* = 0.95 Hz; ipsilateral: *M* = 15.33, *SD* = 0.96 Hz), but in adults, contralateral bursts had a lower peak frequency than ipsilateral bursts (*Z* = -7.05, *p* < 0.001; contralateral: *M* = 21.12, *SD* = 1.77 Hz; ipsilateral: *M* = 21.15, *SD* = 1.78 Hz). There was no difference in frequency span between hemispheres at 9 months (*Z* = -1.13, *p* = 0.258; contralateral: *M* = 2.12, *SD* = 0.83 Hz; ipsilateral: *M* = 2.13, *SD* = 0.82 Hz), but contralateral bursts had a narrower frequency span than ipsilateral bursts at 12 months (*Z* = -5.72, *p* < 0.001; contralateral: *M* = 2.06, *SD* = 0.83 Hz; ipsilateral: *M* = 2.11, *SD* = 0.84 Hz) and in adults (*Z* = -61.38, *p* < 0.001; contralateral: *M* = 1.82, *SD* = 0.76 Hz; ipsilateral: *M* = 2.12, *SD* = 0.83 Hz). Finally, ipsi- and contralateral bursts did not differ in terms of duration at 9 months (*Z* = -1.18, *p* = 0.239; contralateral: *M* = 4.39, *SD* = 2.05 cycles; ipsilateral: *M* = 4.43, *SD* = 2.03 cycles), but contralateral bursts were shorter than ipsilateral bursts in 12-month-olds (*Z* = -5.15, *p* < 0.001; contralateral: *M* = 4.31, *SD* = 2.07 cycles; ipsilateral: *M* = 4.45, *SD* = 2.13 cycles) and in adults (*Z* = -80.19, *p* < 0.001; contralateral: *M* = 4.20, *SD* = 2.42 cycles; ipsilateral: *M* = 5.41, *SD* = 2.80 cycles). Contralateral and ipsilateral beta bursts did not differ in terms of TF-based features at 9 months, but by 12 months, contralateral bursts had a lower peak amplitude, narrower frequency span, and shorter duration than ipsilateral bursts, and by adulthood, contralateral bursts additionally had a lower peak frequency than ipsilateral bursts.

As found in previous studies of movement-related sensorimotor beta activity, changes in the overall burst rate generally tracked changes in mean beta amplitude in the execution condition (Little et al., 2019; Rayson et al., 2022; Figure 3i,j,k). In 9-month-old and 12-month-old infants, there was a bilateral decrease in both the burst rate and mean beta amplitude following the first contact of the hand with the toy (Figure 3i,j). However, in adults, the mean beta amplitude decreased bilaterally following the onset of the movement, whereas the burst rate only decreased contralaterally (Figure 3k). Thus while mean beta amplitude was bilaterally modulated by action execution in all age groups, only changes in the overall beta burst rate were lateralized, and only in adults. Beta activity was only modulated during action observation in adults around the time of the first contact between the hand and toy, and bilaterally for the mean amplitude, whereas the overall burst rate only decreased in C3 (Figure S3).

### Infant beta bursts have the same mean waveform shape as adult bursts

The differences in TF-based beta burst features between infants and adults give little insight into the mechanisms underlying these changes. All of these features are derived from time-frequency decomposition of a time series during a small time window, corresponding to the burst event. Adults typically exhibit beta bursts with a stereotyped wavelet-like shape in the time domain (Bonaiuto et al., 2021; Brady and Bardouille, 2022b; Sherman et al., 2016), which causes the observed features in the TF domain (Jones, 2016; Sherman et al., 2016; Szul et al., 2022). To determine the underlying cause of age-related changes in TF-based features, we compared the waveform shape of beta bursts between 9 and 12-month-olds with adults. The time window for waveform extraction was determined by the FWHM of lagged coherence averaged within the age-specific beta band, yielding 5 cycles (344 ms) for 9-month-olds, 5.2 cycles (341 ms) for 12-month-olds, and 5 cycles (233 ms) for adults. Bursts identified in each age group had a wavelet-like median waveform shape with a strong central negative deflection and surrounding positive deflections (Bonaiuto et al., 2021; Sherman et al., 2016) as well as more peripheral peaks (Brady and Bardouille, 2022b; Szul et al., 2022), but there was great variability around this median (Figure 4a-c). Infant bursts had a greater median waveform amplitude, and because they had a lower peak frequency, had slightly longer waveforms than adult bursts (Figure 4a-c). However, after normalisation and dynamic time warping (Figure S4; Giorgino, 2009), bursts from both infant groups were revealed to have the same qualitative median waveform shape as adult bursts (Figure 4d,e). The normalised distance between the infant burst waveforms and those of adult bursts was slightly lower in 9-month-olds than at 12 months (9m: 0.023, 12m: 0.033). Despite differences in TF-based burst features between age groups, beta bursts have a common median waveform shape across ages, suggesting that they are generated via similar mechanisms that change only quantitatively with age.

**Figure 4.**
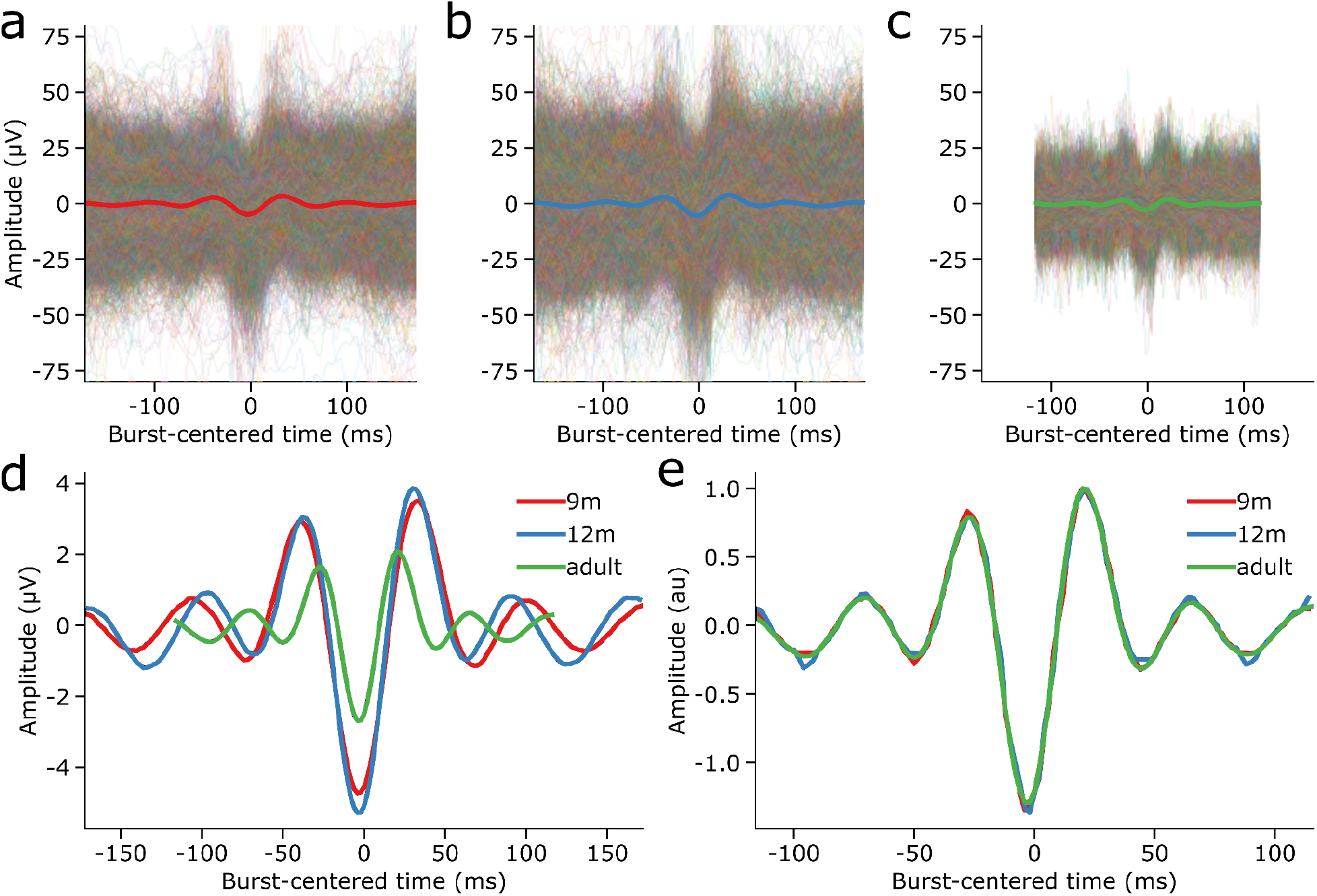
Infant and adult bursts have qualitatively similar burst waveforms. a) The median waveform (thick red line) over all detected beta bursts from 9-month-old participants had a wavelet-like shape, but there was great variability in the waveforms of individual bursts (thin coloured lines). b) As in a, for the 12-month-old participants. c) As in a, for the adult participants. d) The adult beta burst waveforms were, on average, shorter in absolute duration and smaller in amplitude than the mean infant bursts. e) Normalisation and dynamic time warping revealed that the infant and adult beta bursts have qualitatively similar waveform shapes.

### The same burst motifs are increasingly rate-modulated during movement from infancy to adulthood

Although the warped burst waveforms were very similar across age groups, there was a considerable amount of variability around the mean waveform shape. It has been suggested that this variability reflects variability in function (Szul et al., 2022); therefore, we then examined how waveform variability changes with age and thus might drive developmental changes in motor control. We applied principal component analysis (PCA) to the warped, aligned waveforms from all age groups to identify common motifs that explain burst waveform variance across ages. We extracted 14 principal components that explained 80.42% of the waveform variance. To determine which components significantly contributed to waveform variability, we used a permutation test, resulting in 10 significant components (PCs 1-10; *p* < 0.001). Each of these dimensions defined changes in the amplitude or relative amplitude of the central negative deflection, surrounding positive deflections, and peripheral deflections.

We used a single PCA applied globally to all burst waveforms from all ages to compare waveforms between age groups using a common set of dimensions. However, it could be that each component describes waveform variance in a single age group. To verify that the identified burst waveform motifs were not biased towards an overrepresentation of waveform variability from one or two age groups, we ran separate PCAs on bursts from each age group and compared the resulting principal components and burst scores with those from the global PCA applied to all bursts from all age groups. PCs 1-8 were highly similar across ages, both in terms of eigenvectors (9m: *r* = 0.73 - 1.0; 12m: *r* = 0.79 - 1.0; adult *r* = 0.71 - 1.0; Figure S4a-c), and burst scores (9m: *r* = 0.72 - 1.0; 12m *r* = 0.84 - 1.0; adult: *r* = 0.73 - 1.0; Figure S4d-i). For PCs 1-6, the most similar component from each group PCA was the corresponding one of the global PCA, indicating that for these components, not only was the waveform motif very similar, but the relative percentage of variance explained was the same across ages. PCs 7 and 8, however, were reversed in order for the adult PCA, meaning that these motifs explained different amounts of variance in the infant versus adult waveforms. Two of the significant components, PCs 9 and 10, mainly represented waveform variability in the adult participants (PC 9: 9m eigenvector *r* = 0.75, burst score *r* = 0.74; 12m eigenvector *r* = 0.60, burst score *r* = 0.67; adult eigenvector *r* = 0.95, burst score *r* = 0.94; PC 10: 9m eigenvector *r* = 0.57, burst score *r* = 0.61; 12m eigenvector *r* = 0.48, burst score *r* = 0.51; adult eigenvector *r* = 0.72, burst score *r* = 0.74). All but two of the significant burst waveform motifs were therefore common among infants of 9 and 12 months, as well as adults.

We then further analysed four components based on significant changes in the mean burst score in the action execution epochs (no components were significantly modulated in the action observation condition), thus selecting dimensions along which the mean burst shape varied systematically over the course of the trial. Each of these components defined dimensions along which the waveform shape varied markedly from the median waveform (Figure 5a,c,e,g; Figure 6a,c,e,g; see Figures S6-7 for the remaining six significant components). In each of these dimensions, the amplitude of peaks surrounding the central negative deflection, and that of the central deflection itself varied, but the most striking feature of each of the four components is that they represent waveforms with additional peripheral peaks and waveform asymmetry. PCs 3 and 4 defined motifs in which the asymmetry between the surrounding positive deflections and the magnitude of the central negative deflection varied (Figure 5a,c), and PCs 6 and 9 mainly represented changes in the amplitude of the deflections. The mean burst waveform score for each of these components decreased just before or after toy contact, and further decreased during the movement. Hence, not only did the overall burst rate decrease during the movement, but the mean burst waveform shape systematically changed throughout the task.

**Figure 5.**
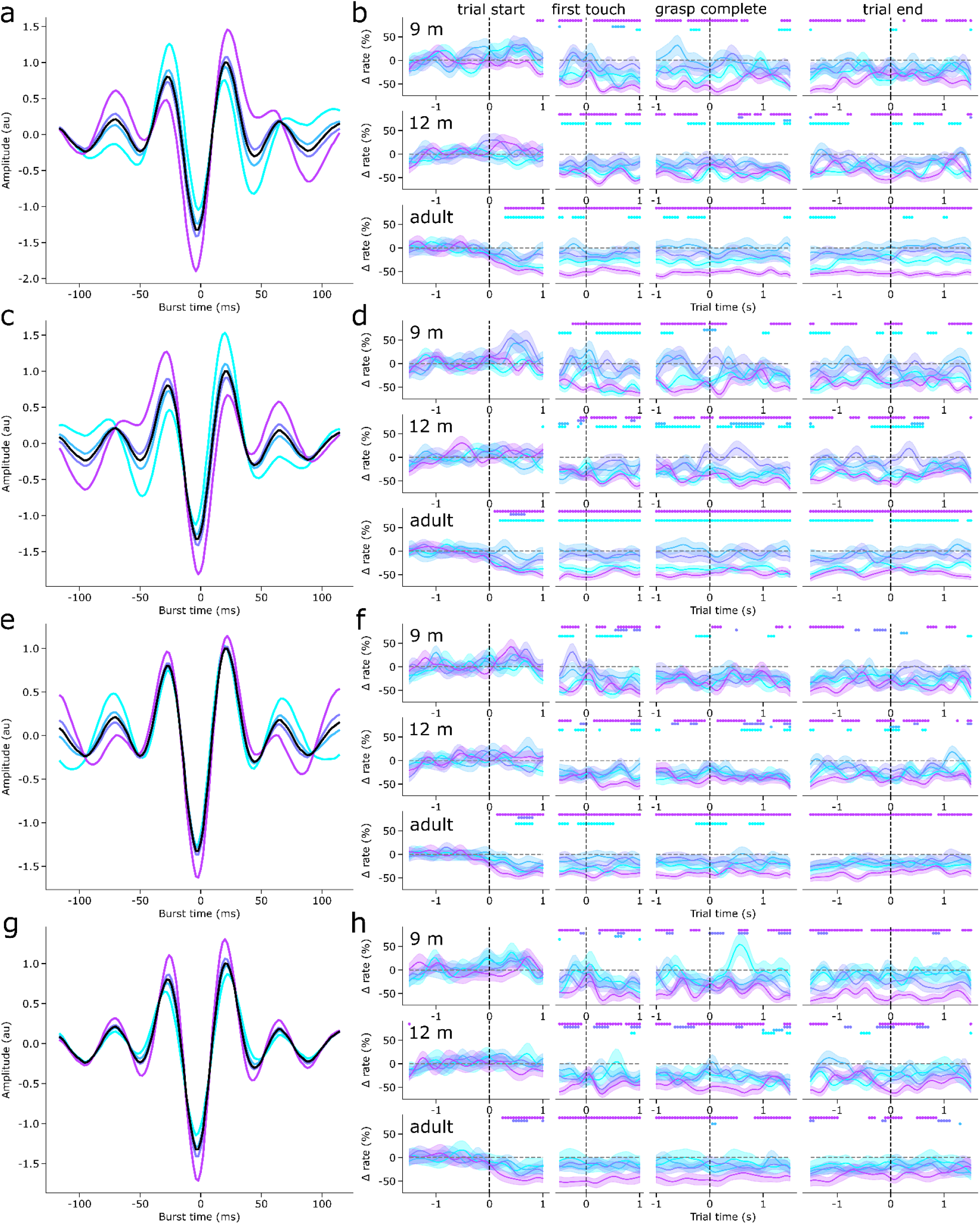
Contralateral burst waveform motifs are increasingly rate-modulated during movement from infancy to adulthood. a) The mean normalised and warped waveforms of beta bursts with scores in four quartiles of PC 3 scores (coloured lines) and the mean overall burst waveform (black). b) The mean baseline-corrected rate of bursts with scores in each PC 3 score quartile (coloured lines, the shaded area indicates the SEM) over the course of the (columns, from left to right) trial start, first touch, grasp completion, and trial end epochs in the contralateral centre cluster for 9-month-old (top row), 12-month-old (middle row), and adult (bottom row) participants. The coloured dots indicate where the burst rate in the corresponding score quartile is different from baseline. c-d) As in a-b, for PC 4. e-f) As in a-b, for PC 6. g-h) As in a-b, for PC 9.

**Figure 6.**
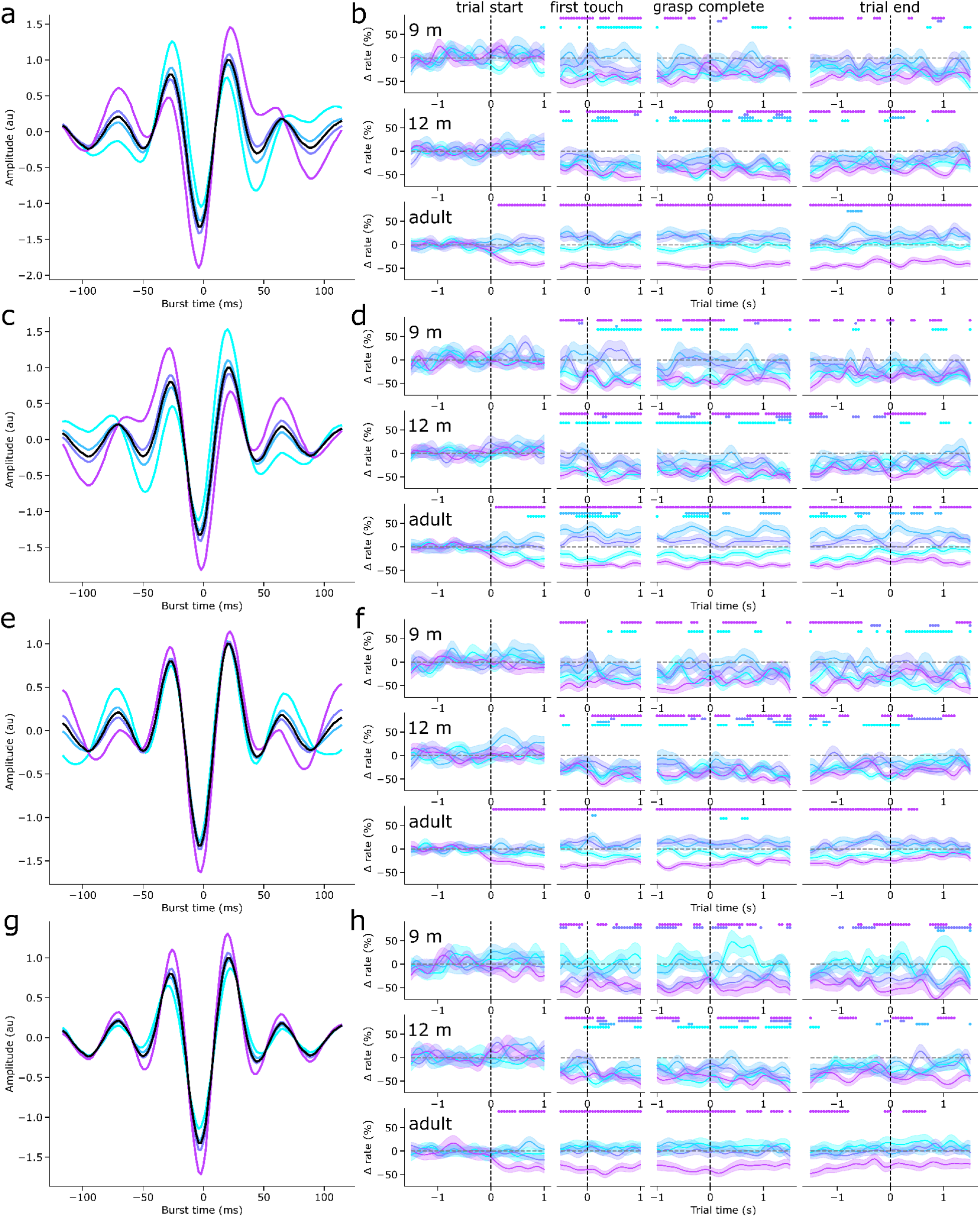
Ipsilateral beta burst motifs rate-modulated differently from contralateral motifs during movement in adulthood, but not infancy. a) The mean normalised and warped waveforms of beta bursts in with scores in four quartiles of PC 3 scores (coloured lines) and the mean overall burst waveform (black). b) The mean baseline-corrected rate of bursts with scores in each PC 3 score quartile (coloured lines, the shaded area indicates the SEM) over the course of the (columns, from left to right) trial start, first touch, grasp completion, and trial end epochs in the ipsilateral central cluster for 9-month-old (top row), 12-month-old (middle row), and adult (bottom row) participants. The coloured dots indicate where the burst rate in the corresponding score quartile is different from baseline. c-d) As in a-b, for PC 4. e-f) As in a-b, for PC 6. g-h) As in a-b, for PC 9.

The systematic changes in overall burst rate and mean waveform shape do not indicate whether there was an increase in bursts with low component scores, or a decrease in bursts with high component scores. To investigate whether different waveform motifs were differentially rate-modulated during movement, we therefore examined the burst rate according to the waveform shape in the execution condition. For each of the four selected principal components, we binned bursts into four quartiles based on their score along that dimension, as well as the time during the trial in which they occurred. We then computed the burst rate for each PC score quartile, baseline-corrected the burst rates, and used permutation tests to determine significant deviations from the baseline. Along PCs 3, 4, and 6, bursts with scores in the first and fourth quartiles decreased in rate contralaterally during the movement compared to baseline, whereas those in the second and third quartiles (with waveform shapes closer to the median waveform) were not rate-modulated (Figure 5b,d,f). For PC 9, only bursts in the fourth quartile decreased in rate (Figure 5h). Interestingly, the decrease in rate of bursts in the first and fourth quartiles occurred just after the onset of the trial in adults, and only just before or after the moment the hand touched the toy in 9-month and 12-month-olds. Moreover, this decrease was significantly different from the baseline rate over longer periods of time in adults than in infants. Ipsilaterally, in adults, for the most part only bursts in the fourth quartiles of PCs 3, 4, 6, and 9 decreased in rate during movement (Figure 6), and for PC 4, bursts closer to the median waveform actually increased in rate during movement (Figure 6d). In infants, the rate of ipsilateral bursts in the first and fourth quartiles of each selected PC decreased during movement, similar to contralateral bursts. The other significant components were not reliably differentially rate-modulated during action execution in any age group, or only in adults (Figures S6, S7). Beta bursts therefore occur within specific burst waveform motifs, some of which reduce in rate increasingly earlier during movement from infancy to adulthood, and others which are differentially rate-modulated ipsilateral to movement only in adulthood.

## Discussion

The current study provides insights into the developmental trajectory of beta bursts and their potential role in sensorimotor processing. Our findings demonstrate that infant beta activity, similar to adults, manifests as transient bursts with diverse spectral and temporal features, albeit at a lower peak frequency. The lateralization of beta burst features increases between 9 months and 12 months, and further into adulthood. We observed that infant beta bursts exhibit a mean waveform shape similar to adults and occur within a set of common motifs at all ages, indicating a possible shared mechanism of generation. Interestingly, several motifs classified beta bursts into those that decrease in rate during movement and those whose rate does not change or slightly increases, suggesting that different burst types may be involved in distinct sensorimotor processes. Moreover, our results demonstrate that the bursts with waveforms furthest from the mean waveform decrease more in rate and earlier during movement with age. In adults, the burst rate of certain motifs is differentially modulated ipsi- and contralateral to the hand used for grasping, but infant beta bursts in these same motifs are bilaterally rate-modulated during movement. Taken together, these findings highlight the complexity and diversity of beta bursts and their potential role in sensorimotor development.

The fact that infant and adult beta bursts have qualitatively similar mean waveform shapes suggests that they are generated by the same mechanism, which changes in a predominantly quantitative way with age. The dominant computational model of beta burst generation proposes that beta bursts are driven by temporally aligned synaptic inputs to deep and superficial layers that drive intracellular current in opposite directions within a cortical column (Law et al., 2022; Sherman et al., 2016). The combination of these two currents results in a net cumulative dipole with the same waveform as the mean burst waveforms we observed. This model was based on bursts observed in the somatosensory cortex, but experimental support for the model has been found in the human motor cortex (Bonaiuto et al., 2021; Szul et al., 2022). It has been suggested that in the somatosensory cortex, these two inputs arise from the lemniscal and non-lemniscal thalamus (Sherman et al., 2016). However, this model is unable to generate bursts with the rich variety of waveforms observed in the human motor cortex (Szul et al., 2022), suggesting that a more complex pattern of inputs might generate beta bursts in this region, which is in line with its wider afferent connectivity profile (Geng et al., 2022; Hooks, 2017). The primary motor cortex receives projections from different thalamic motor nuclei that relay information from the basal ganglia and cerebellum and target predominantly to the deep and superficial layers, respectively (Hooks et al., 2013; Kuramoto et al., 2015, 2009), as well as from thalamic sensory nuclei that mainly target the superficial layers (Hooks et al., 2013; Kuramoto et al., 2015, 2009; Ohno et al., 2012). These projections may account for the stereotypical mean burst waveform shape, but the motor cortex also receives lamina-specific inputs from the sensory and frontal cortices (Hira et al., 2013; Hooks et al., 2013; Mao et al., 2011; Rouiller et al., 1993), including six premotor areas (Dum and Strick, 2002). This confluence of synaptic inputs may contribute to the observed variability of burst waveforms in a context-dependent way (Szul et al., 2022). Myelination of inter-regional fibre tracts is particularly intensive during the first year of life (Dubois et al., 2014), which shapes neurophysiological activity measured with EEG (Adibpour et al., 2017), and therefore maturation of thalamo-cortical and cortico-cortical projections may explain differences in the peak frequency of infant and adult beta bursts.

Our results suggest that different types of beta bursts are involved in different sensorimotor mechanisms. Interestingly, we observed that bursts with certain waveforms decreased in rate earlier during movement in adults compared to infants. This may be due to the increasing use of visual information for movement preparation and planning, and more generally, a shift from reflexive to prospective control of movement (Hofsten, 1993; van der Meer et al., 1994; Witherington, 2005). Newborn infants exhibit basic arm movements towards fixated objects (von Hofsten, 1982), which are essential precursors to object-directed reaching and grasping (Bhat et al., 2005; Bhat and Galloway, 2006). As grasping develops throughout infancy and early childhood, there is an increasing reliance on visual information to preprogram the grasp (Lasky, 1977), and infants begin to pre-orient their wrists (Lockman et al., 1984; McCarty et al., 2001) and shape their hands (Fagard, 2000; Fagard and Jacquet, 1996; von Hofsten and Rönnqvist, 1988; Witherington, 2005) to match the object’s position, orientation, and shape. Computational models of this learning process suggest that these motor programs are learned via trial-and-error reinforcement learning (Bonaiuto and Arbib, 2015; Oztop et al., 2004), which shapes projections from object representations in the parietal cortex to grasp motor program representations in the premotor cortex, thereby altering the information that reaches the motor cortex. It may be that certain burst types represent the activation of motor representations (Little et al., 2019), and at rest, constant micro-movements, or “non-task-related movements” (Musall et al., 2019; Nougaret et al., 2023; Stringer et al., 2019) are associated with the occurrence of beta bursts in specific motifs. Once a motor representation for a selected movement has been activated, these extraneous movements may be reduced (D’Souza et al., 2017) while the movement is prepared, resulting in a decrease in burst rate. However, in this study, adults performed faster reaching movements than infants, and therefore the earlier burst rate decrease we observed in adults may simply reflect the fact that their hand was closer to the toy at the same time post trial start. Future studies should therefore track movement kinematics in a more detailed way in order to account for this difference.

Consistent with our behavioural results, young infants exhibit an ambidextrous pattern of hand usage, without any distinct preference for either hand (Corbetta and Thelen, 1999; Rönnqvist and Domellöf, 2006). However, a bias for using one hand emerges by about 1 year (Hinojosa et al., 2003), develops into a clear preference by approximately 18 months of age, (Fagard and Marks, 2000), and subsequently becomes more pronounced in early childhood (Ingram, 1975). Whilst we observed a change in burst feature lateralization between 9 and 12 months, the overall burst rate was only lateralized in adults. Sensorimotor activity is not hemispherically lateralized in neonates (Erberich et al., 2006), which may be why there were no hemispheric differences in either burst features or burst rate at 9 months. The increasing lateralization of beta burst activity between 9 months and adulthood may be due to the decrease in interhemispheric functional connectivity between motor cortices that occurs during the first year of life (Xiao et al., 2018), and the ongoing development of interhemispheric inhibition in childhood (Ciechanski et al., 2017). However, the pattern of activity that we observed in adults has implications for the functional role of beta activity. If beta activity is purely inhibitory in nature (Picazio et al., 2014; Zhang et al., 2008), one would expect either a decrease in contralateral beta activity during movement, thus disinhibiting the limb to be moved, and either no change or an increase in ipsilateral beta activity. Consistent with this view, we observed a decrease in overall burst rate contralaterally in adults, and no change in overall ipsilateral burst rate. However, we found four main types of burst rate lateralization: bursts with a static rate bilaterally, bursts that remained static contralaterally and increased in rate ipsilaterally, bursts that decreased in rate contralaterally and were static ipsilaterally, and bursts that decreased in rate bilaterally. These findings suggest that the role of beta activity in sensorimotor processes is more complex than a simple inhibitory effect.

Burst rate and beta power were only modulated during action observation in adults, but there were no consistent differences between age groups in the kinematics of the observed movement. Previous work has found a reduction in mu activity during action observation in infants and adults (Cannon et al., 2014; Filippi et al., 2016; Marshall et al., 2011; Marshall and Meltzoff, 2011), and beta activity in adults (Press et al., 2011), but beta activity has only been shown to be modulated during action observation in older infants (van Elk et al., 2008). Motor simulation theory suggests that observation of someone else performing an action activates an internal sensorimotor simulation of performing that same action (Jeannerod, 2001; Miall, 2003; Press et al., 2011; Wolpert et al., 2003). If certain burst types are related to the activation of internal models, one possible explanation for this finding is that these internal models are not yet well developed in infants and are therefore not activated during action observation. This is in line with the increasing amount and earlier timing of rate modulation of these burst motifs from infancy to adulthood during action execution. However, even in adults, beta modulation during action observation was mostly confined to the period from the start of the trial to the completion of the grasp, and it returned to baseline levels during the manipulation of the toy. It is possible that the action used in the observation condition here, grasping with a claw-like tool, was insufficiently familiar both to the infants and adults, and the beta modulation we observed in adults was actually due to the reach and grasp of the tool, rather than the use of the tool to grasp the toy. However, this would require more precise tracking of the hand kinematics and a comparison with a simple grasping action.

In line with the few previous studies that have identified age-specific beta frequency bands in infancy (Meyer et al., 2016; Rayson et al., 2022), we found the peak beta frequency to be around 15 Hz in 9- and 12-month-old infants. One factor that may have contributed to the lack of previous research on infant beta activity is the fact that in infants, muscle artefacts from jaw and arm movements appear at 15 Hz in peripheral electrodes, spectrally overlapping the sensorimotor beta frequency range (Georgieva et al., 2020). However, it is unlikely that our results are driven by these artefacts because the spatial topography of lagged coherence in the identified beta band localises very strongly to the central electrodes, and we found a decrease in burst rate during movement, when one would expect an increase in muscle artefacts. The transient nature of bursting activity in any frequency range means that burst detection methods based on power thresholds are prone to erroneously identify any artefact whose waveform yields beta power when the Fourier transform is applied. These artefacts would also be included in any analysis of frequency band-specific average power. The approach used here identifies all potential burst events in TF space and then allows them to be sorted and analysed according to their waveform shape, a less biassed reflection of the underlying neural activity.

The present study has several limitations worth noting. Our results suggest developmental shifts in the strength, timing, and duration of different synaptic drives to the sensorimotor cortex, thus modulating the features and rate of sensorimotor beta bursts. However, without any measure of anatomical or functional connectivity, it is impossible to know where these projections originate from. This could be determined using measures of white matter microstructure in fibre tracts or resting state functional connectivity metrics to determine which projections to sensorimotor cortex modulate burst waveforms. In this study, the actual three-dimensional trajectories of the arm and hand during performed and observed movements were not quantified, limiting the granularity of the conclusions that can be drawn about the relationship between beta activity and sensorimotor processes. Future studies could take advantage of recently developed techniques for markerless behavioural tracking (Desmarais et al., 2021; Karashchuk et al., 2021; Mathis et al., 2018), which is especially important with infants as previous motion tracking approaches require markers to be attached to the body. Finally, although this study included multiple age groups, it was cross-sectional. The ideal future study for determining the developmental role of beta bursts would therefore be longitudinal from infancy to early childhood, and involve MRI, M/EEG, and markerless kinematic tracking to determine the precise relationship between afferent brain regions driving beta bursts, burst characteristics, and movement.

Several developmental disorders such as attention deficit/hyperactivity disorder (in a subset of children; Clarke et al., 2013, 2007) and autism spectrum disorder (Coben et al., 2008; Tierney et al., 2012) are associated with aberrant beta activity. Altered sensorimotor beta activity is also seen in the context of cerebral palsy and other movement disorders linked to early brain injury and/or atypical neurodevelopment (Démas et al., 2020), though studies with infants and young children are lacking. Early diagnosis is crucial for identifying individuals who would benefit from therapeutic intervention (Landa, 2008; Rommel et al., 2015), with the most effective interventions implemented during sensitive periods of brain development. However such interventions require sensitive biomarkers to determine their efficacy in their early stages because their behavioural effects may be delayed (Cioni et al., 2016; Finlay-Jones et al., 2019; Morris et al., 2023). Burst waveform shape could therefore provide the required sensitivity by allowing detection of abnormal burst waveforms, altered rate-modulation of certain types of bursts, or atypical lateralization of burst features or rate, all of which would be masked by nonspecific measures of highly averaged beta power. More generally, comparison of beta burst activity in typical versus atypical motor development trajectories may be instrumental in teasing apart the mechanistic functional role of normative beta activity.

In summary, this study provides novel insights into the nature of beta bursts in both infant and adult populations, shedding light on their shared and distinct characteristics. Specifically, our results demonstrate that beta activity in both groups is largely composed of bursts with similar waveform shapes, suggesting a shared mechanism of generation. However, we found important differences between infants and adults, including lower peak frequency, later decrease in burst rate during movement, and bilaterally rate-modulated bursts in infants, as compared to differential modulation in adults. These findings suggest that variability in burst characteristics may be indicative of variability in sensorimotor processes, and that specific burst types may undergo changes in their rate modulation throughout development. Overall, our study provides new insights into the developmental trajectory of beta bursts, and highlights their potential role as a fundamental aspect of the sensorimotor system during early development.

## Acknowledgements

This research was supported by a grant from the National Institute of Health (NIH) (P01 HD064653) to NAF and PFF. HR, MJS, PEK, MGM, and JJB were supported by the European Research Council (ERC) under the European Union’s Horizon 2020 research and innovation programme (ERC consolidator grant 864550 to JJB). The funders had no role in the preparation of the manuscript.

## Supplementary Information

**Table S1.** Participants and trials used in analyses. The number of subjects per age group with at least 5 trials, and the range, mean, and standard deviation of the number of trials per subject for each epoch.

|  |  | 9m |  | 12m |  | adults |  |
| --- | --- | --- | --- | --- | --- | --- | --- |
|  |  | N | Trials | N | Trials | N | Trials |
| Execution | Start | 27 | 5-19, M = 13.00, SD = 3.38 | 31 | 6-20, M = 12.55, SD = 4.75 | 22 | 11-20, M = 19.50, SD = 1.92 |
|  | Touch | 22 | 5-18, M = 10.09, SD = 3.26 | 25 | 5-18, M = 10.36, SD = 3.51 | 22 | 10-20, M = 19.45, SD = 2.13 |
|  | Grasp | 22 | 5-18, M = 10.14, SD = 3.41 | 25 | 5-18, M = 10.40, SD = 3.45 | 22 | 10-20, M = 19.45, SD = 2.13 |
|  | End | 22 | 5-17, M = 9.82, SD = 3.19 | 24 | 5-17, M = 10.12, SD = 3.48 | 22 | 10-19, M = 18.50, SD = 1.92 |
| Observation | Start | 28 | 5-19, M = 13.57, SD = 4.34 | 32 | 5-19, M = 12.72, SD = 4.31 | 22 | 11-20, M = 18.68, SD = 1.73 |
|  | Touch | 28 | 5-19, M = 13.79, SD = 4.40 | 33 | 5-19, M = 12.85, SD = 4.54 | 22 | 11-20, M = 18.68, SD = 1.73 |
|  | Grasp | 28 | 5-19, M = 13.79, SD = 4.39 | 33 | 5-19, M = 12.91, SD = 4.56 | 22 | 11-20, M = 18.68, SD = 1.73 |
|  | End | 27 | 8-19, M = 13.74, SD = 4.05 | 33 | 5-19, M = 12.55, SD = 4.44 | 22 | 10-20, M = 18.64, SD = 1.94 |

**Figure S1.**
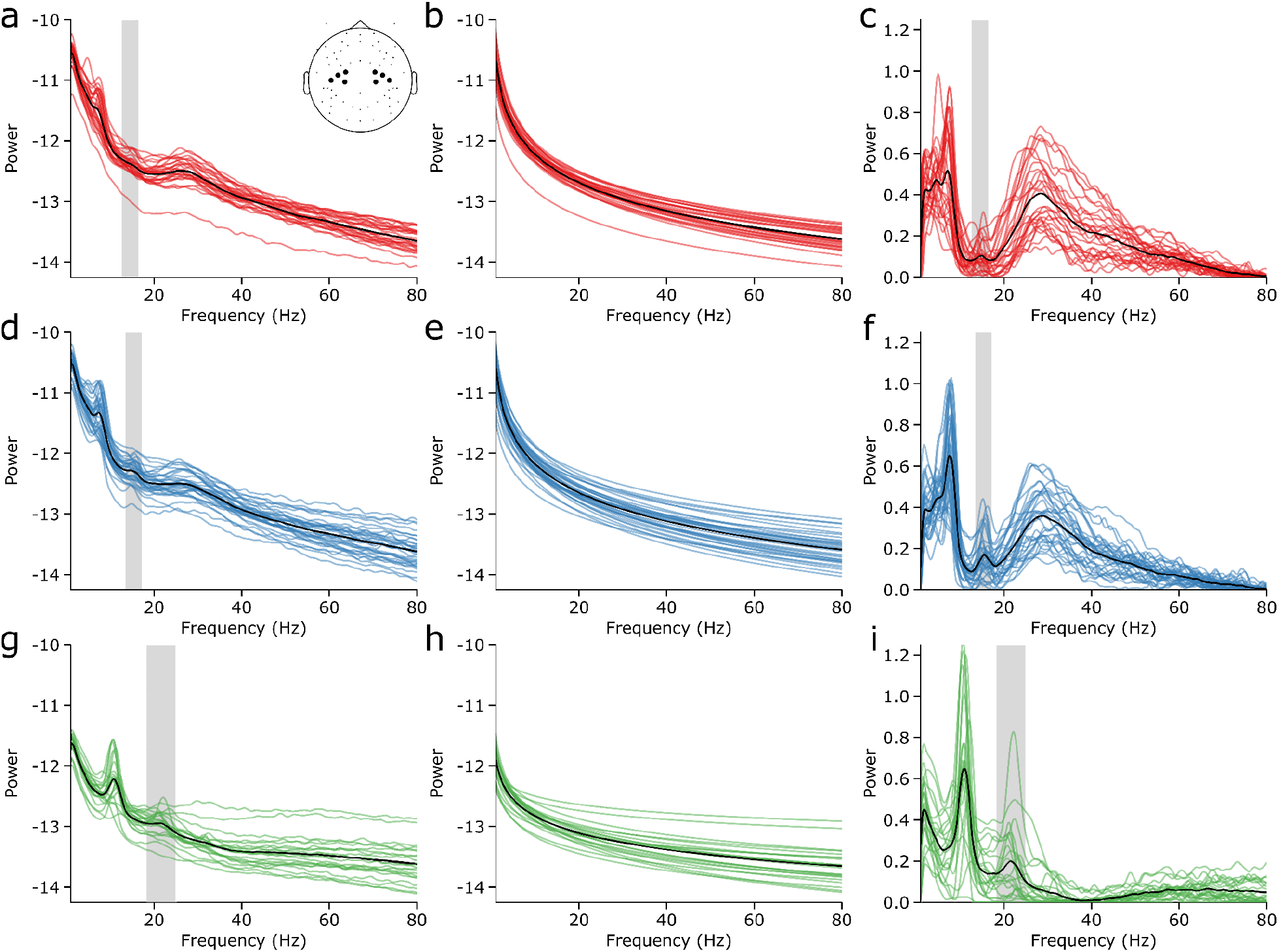
Individual participant power spectra in the C3 and C4 clusters. a-c) PSDs (a), estimated aperiodic components (b), and residual PSDs (c) from the C3 and C4 clusters for each 9-month-old participant. The grey shaded region indicates the limits of the identified beta band. d-f) As in a-c, for 12-month-old participants. g-i) As in a-c, for adult participants.

**Figure S2.**
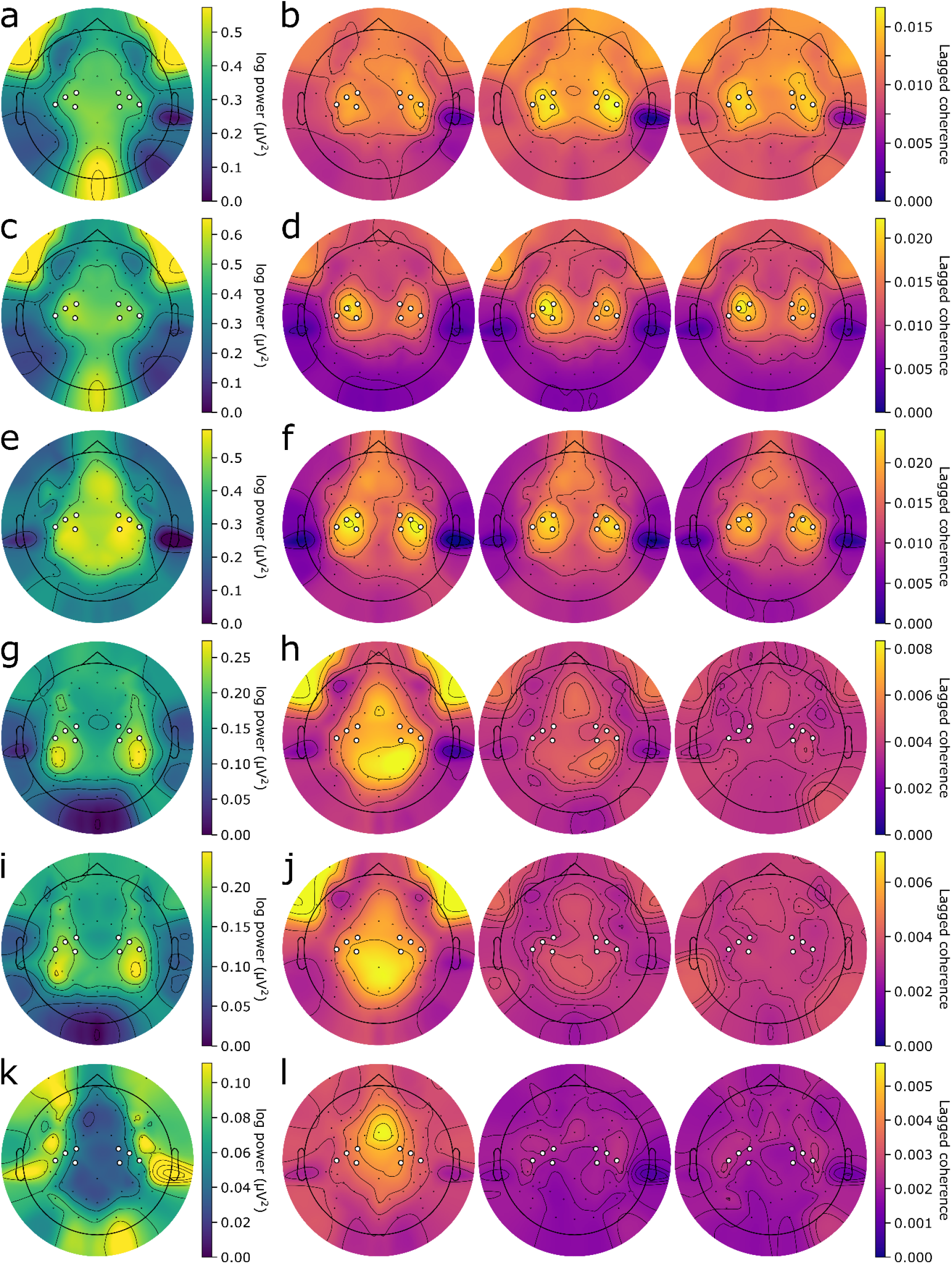
Power and lagged coherence topographies for alpha/mu and high frequency activity. a) Topography of periodic power in the alpha/mu band, averaged over 9-month-old participants. b) Alpha/mu lagged coherence topographies at (from left to right) 2, 3, and 4 cycles, averaged over 9-month-old participants. c-d) As in a-b, for the 12-month-old participants. e-f) As in a-b, for the adult participants. g-h) As in a-b, for high frequency (> 40 Hz) activity. i-j) As in g-h, for the 12-month-old participants. k-l) As in g-h, for the adult participants.

**Figure S3.**
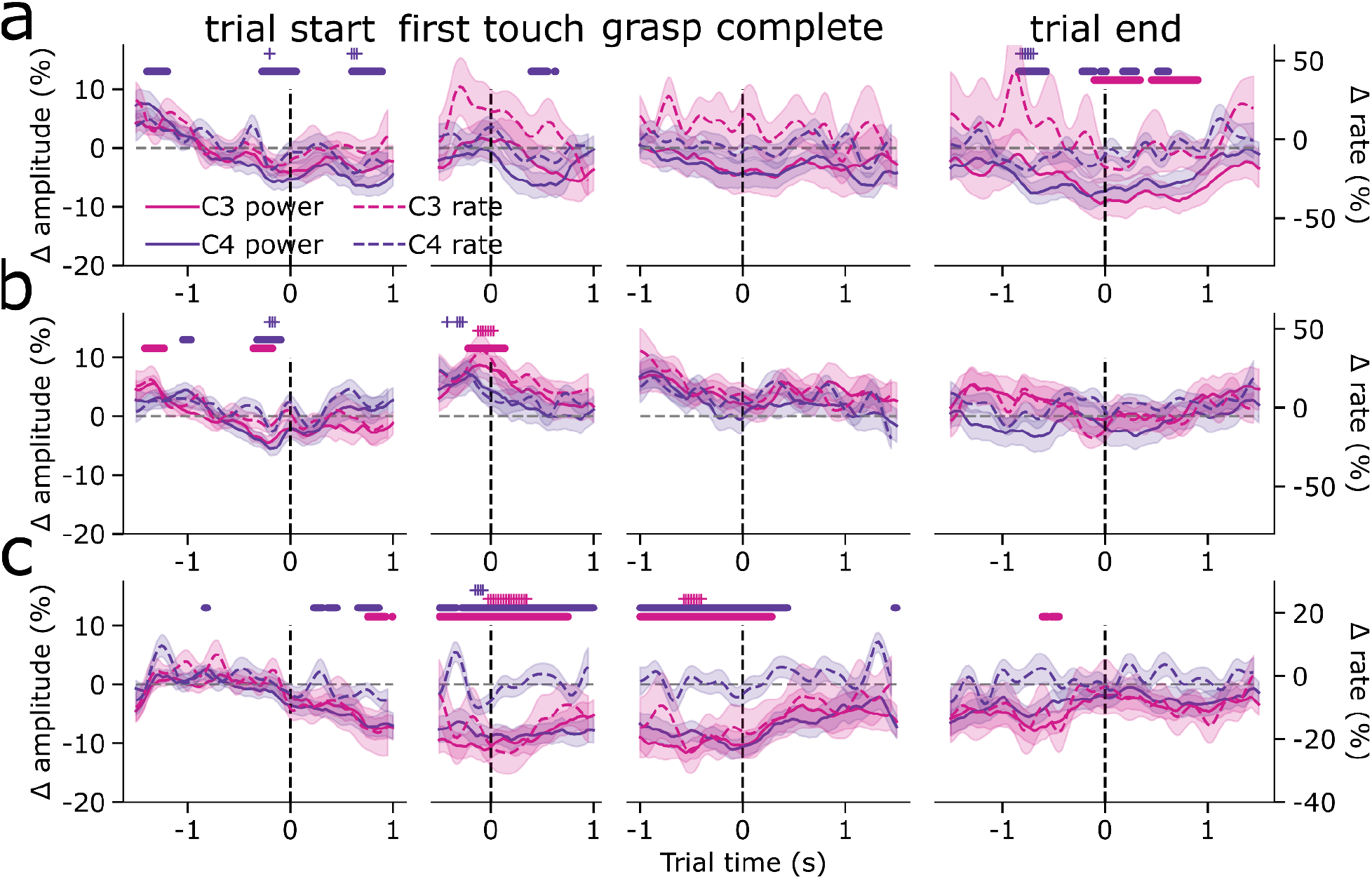
Beta activity is only modulated during action observation in adulthood. a) The mean burst rate (dashed lines; shaded area shows the standard error) and mean beta amplitude (solid lines; shaded area shows the standard error) in the C3 and C4 clusters for the 9-month-old participants in the observation condition. The coloured dots and asterisks indicate times in which beta amplitude or burst rate significantly deviated from baseline. b) As in a, for the 12-month-old participants. c) As in a, for the adult participants.

**Figure S4.**
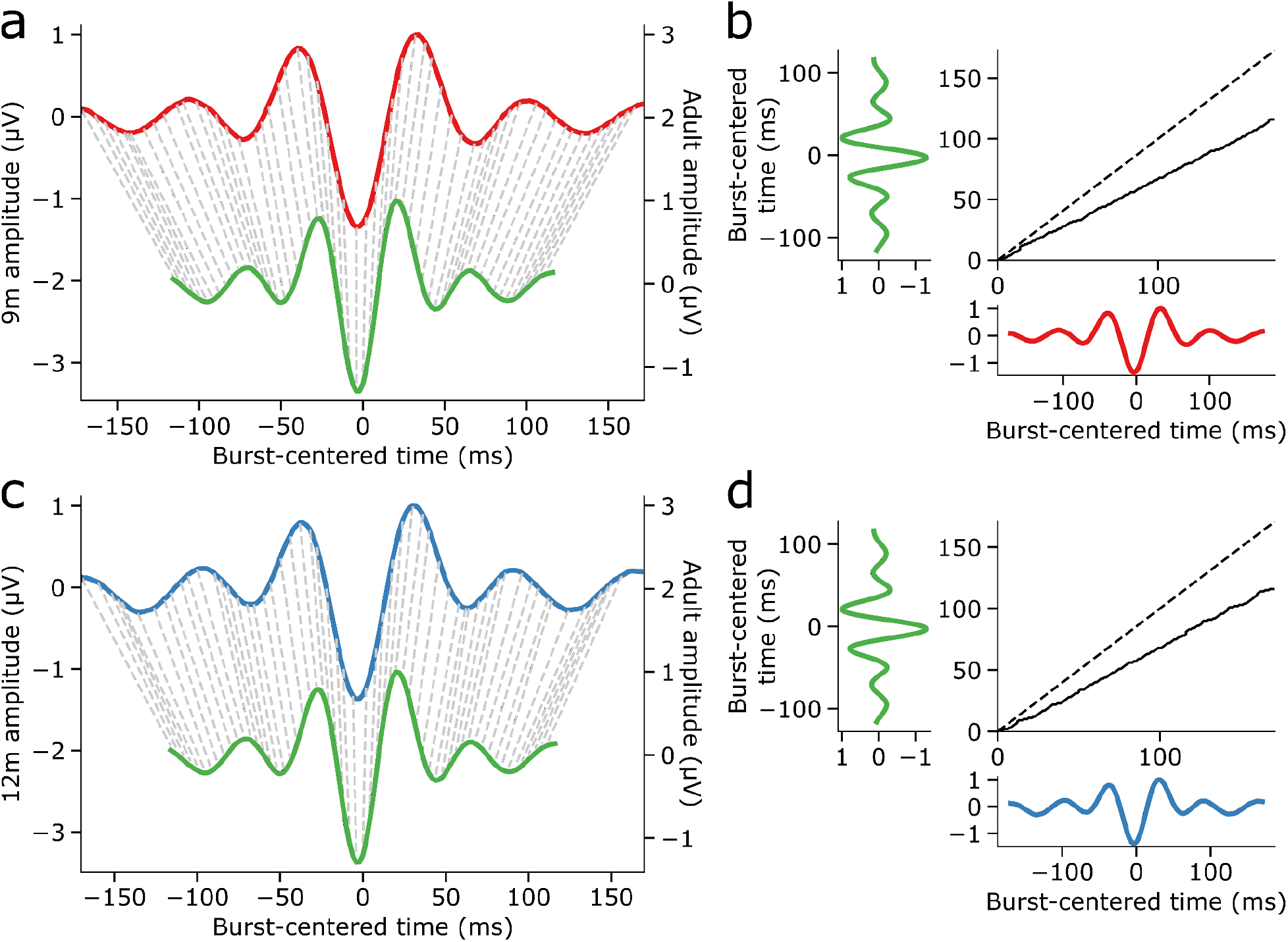
Dynamic time warping of infant burst waveforms to adult bursts. a) Correspondence between time points (dashed lines) of 9-month-old beta burst waveforms (red) and adult beta burst waveforms (green). b) The alignment curve (black solid line) resulting from dynamic time warping of beta burst waveforms from 9-month-old participants to those of adults. The dashed line indicates the alignment curve for two already aligned signals. c-d) As in a-b, for 12-month-old participants.

**Figure S5.**
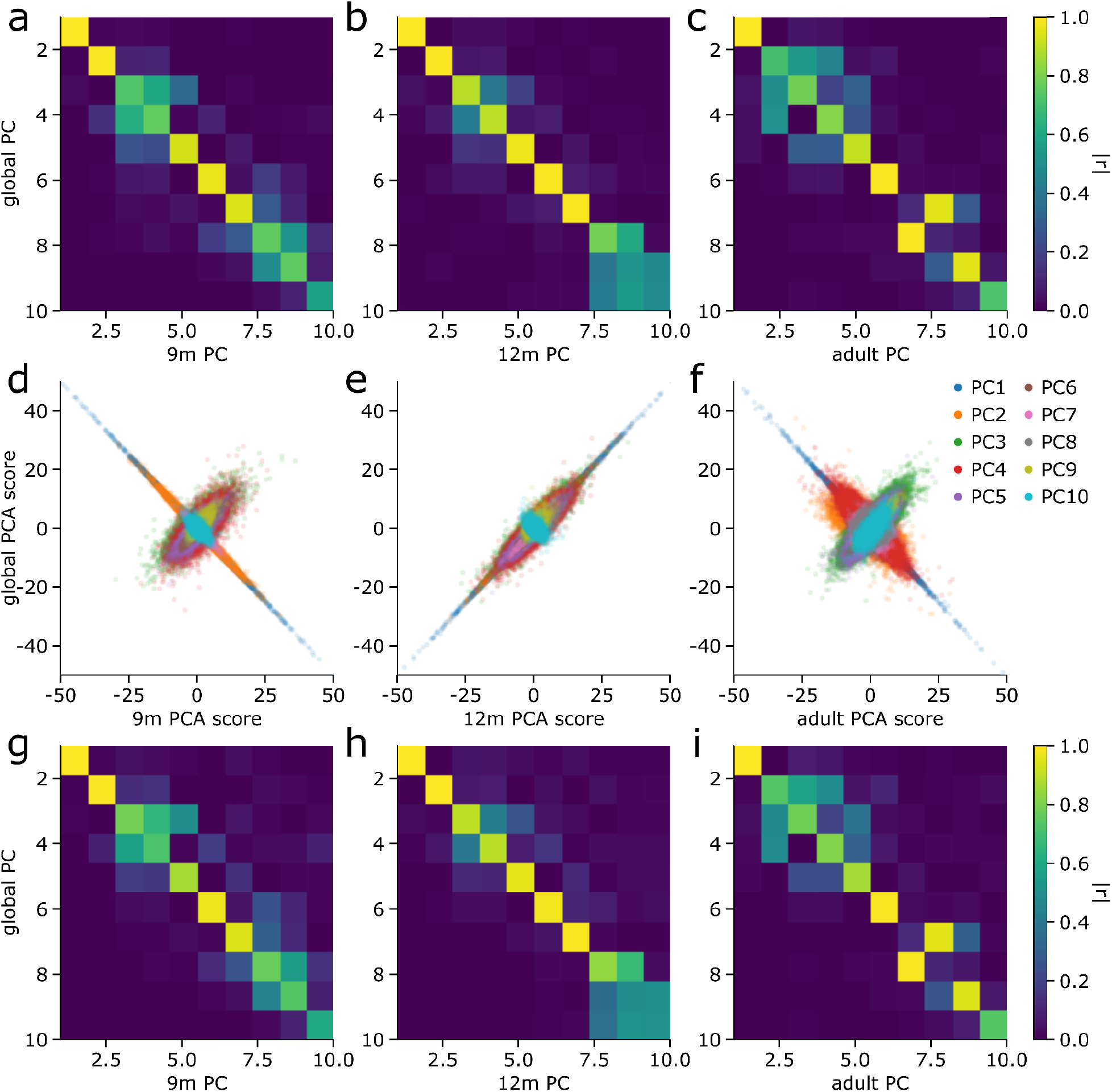
Global versus age group-specific PCA. a) Absolute value of the correlation coefficient between eigenvectors from the PCA ran only on 9-month-old bursts, and those from the global PCA. b-c) As in a, for 12-month-olds (b), and adults (c). d) For each burst detected in 9-month-olds, the score for each PC from PCA ran on only the 9-month-old bursts (x-axis) versus the score from the PCA ran on all bursts over all ages. e-f) As in d, for 12-month-olds (e), and adult participants (f). g) Absolute value of Pearson’s correlation coefficient between scores for 9-month-old infant bursts from PCA applied only to 9-month-old bursts and the global PCA, for each PC. h-i) As in g, for 12-month-olds (h), and adult participants (i).

**Figure S6.**
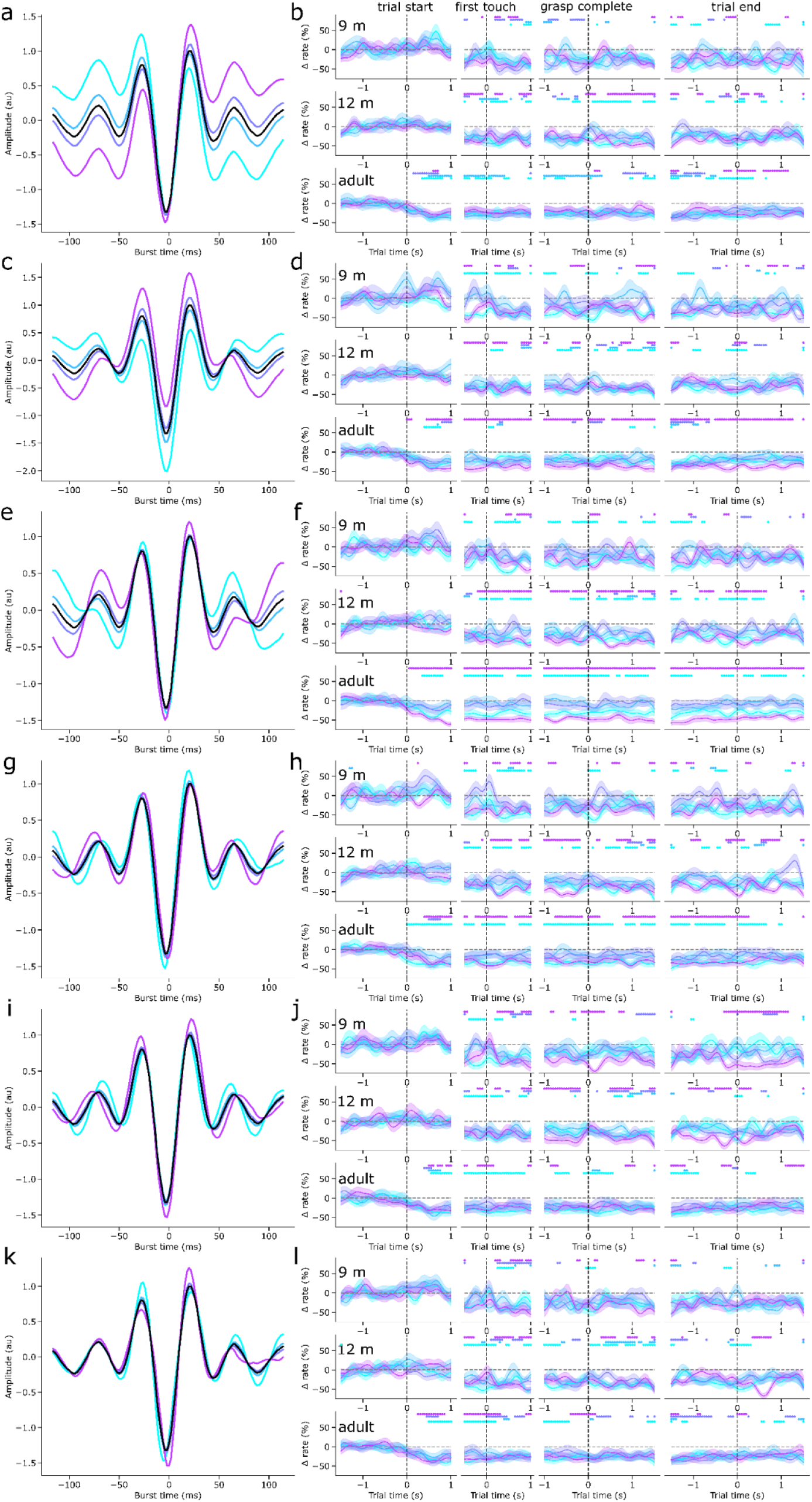
Contralateral beta burst motifs. a) The mean normalised and warped waveforms of beta bursts with scores in four quartiles of PC 1 scores (coloured lines) and the mean overall burst waveform (black). b) The mean baseline-corrected rate of bursts with scores in each PC 1 score quartile (coloured lines, the shaded area indicates the SEM) over the course of the (columns, from left to right) trial start, first touch, grasp completion, and trial end epochs in the contralateral centre cluster for 9-month-old (top row), 12-month-old (middle row), and adult (bottom row) participants. The coloured dots indicate where the burst rate in the corresponding score quartile is different from baseline. c-d) As in a-c, for PC 2. e-f) As in a-b, for PC 5. g-h) As in a-b, for PC 7. i-j) As in a-b, for PC 8. k-l) As in a-b, for PC 10.

**Figure S7.**
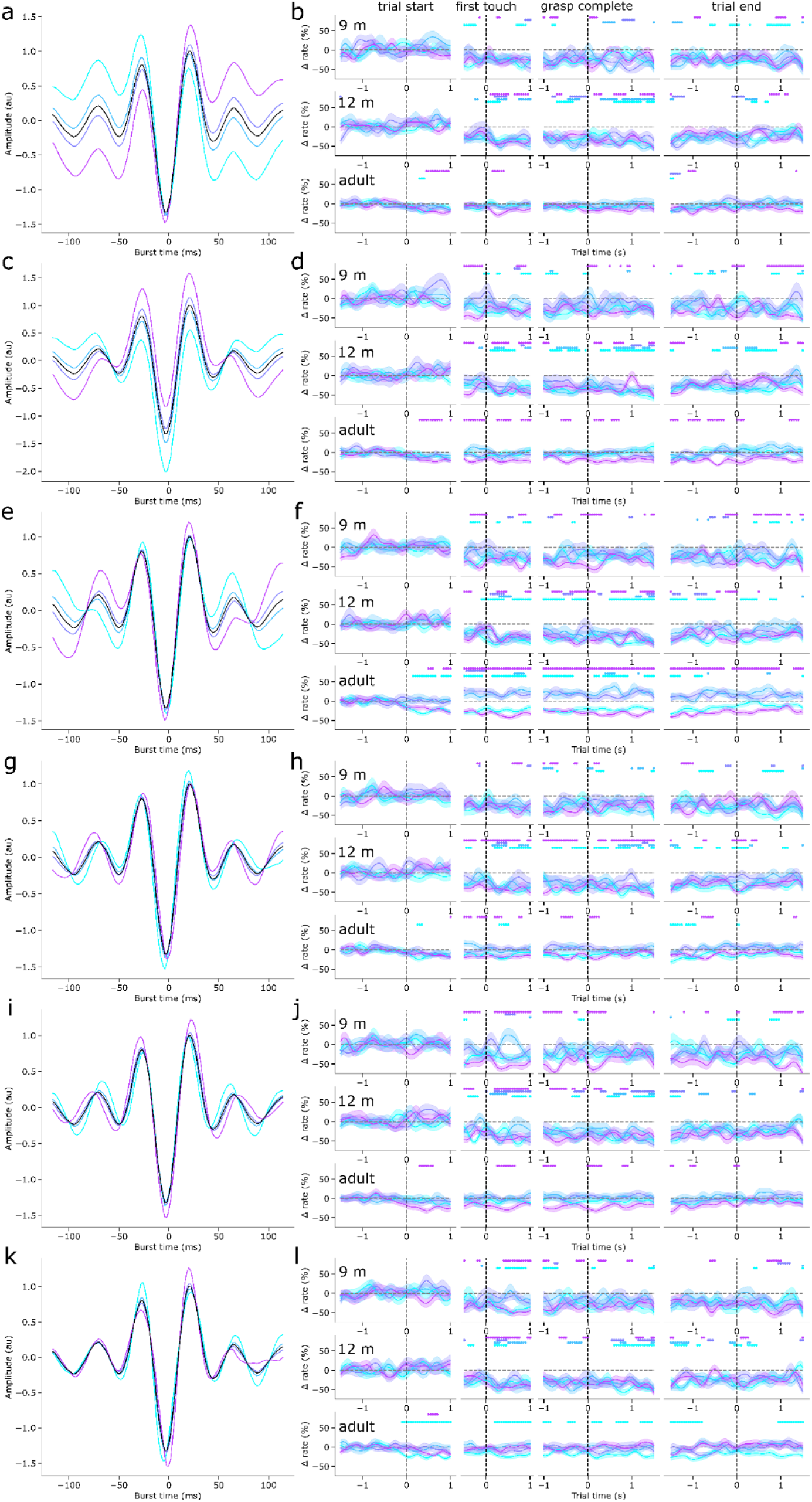
Ipsilateral beta burst motifs. a) The mean normalised and warped waveforms of beta bursts with scores in four quartiles of PC 1 scores (coloured lines) and the mean overall burst waveform (black). b) The mean baseline-corrected rate of bursts with scores in each PC 1 score quartile (coloured lines, the shaded area indicates the SEM) over the course of the (columns, from left to right) trial start, first touch, grasp completion, and trial end epochs in the ipsilateral centre cluster for 9-month-old (top row), 12-month-old (middle row), and adult (bottom row) participants. The coloured dots indicate where the burst rate in the corresponding score quartile is different from baseline. c-d) As in a-c, for PC 2. e-f) As in a-b, for PC 5. g-h) As in a-b, for PC 7. i-j) As in a-b, for PC 8. k-l) As in a-b, for PC 10.

